# MCL-1 regulates cellular transitions during oligodendrocyte development

**DOI:** 10.1101/2024.12.20.629796

**Authors:** Melanie Gil, Marina R Hanna, Vivian Gama

## Abstract

Oligodendrocytes are the myelinating cells of the central nervous system. Regulation of the early stages of oligodendrocyte development is critical to the function of the cell. Specifically, myelin sheath formation is an energetically demanding event that requires precision, as alterations may lead to dysmyelination. Recent work has established that fatty acid β-oxidation is required for the function of oligodendrocytes. We have shown that MCL-1, a well-characterized anti-apoptotic protein, is required for the development of oligodendrocytes *in vivo*. Further, it was recently uncovered that MCL-1 regulates long-chain fatty acid β-oxidation through its interaction with acyl-CoA synthetase long-chain family member 1 (ACSL1), an enzyme responsible for the conversion of long-chain fatty acids into acyl-CoA. Here, we introduce an *in vitro* system to isolate human stem cell-derived oligodendrocyte progenitor cells and investigate the involvement of MCL-1 during human oligodendrocyte development. Using this system, we pharmacologically inhibited MCL-1 in oligodendrocyte progenitor cells (OPCs) to elucidate the non-apoptotic function of the protein at this developmental stage. Additionally, we used a motor neuron co-culture system to investigate the downstream effects that MCL-1 inhibition has at later developmental stages when oligodendrocytes begin to contact axons and generate myelin basic protein. We demonstrate that the mitochondrial network changes in human oligodendrocyte development resemble those reported *in vivo*. Our findings point to MCL-1 as a critical factor essential at the OPC stage for proper oligodendrocyte morphogenesis.

## Introduction

Oligodendrogenesis, the production of myelin-forming oligodendrocytes, is a fundamental process during brain development. This process demands a lot of energy from oligodendrocytes, which require the capacity to generate ATP and lipids to form and maintain the myelin sheath. Additionally, oligodendrocytes provide metabolic support to the axons they myelinate, increasing their bioenergetic demands. Evidence from previous studies demonstrates that oligodendrocyte precursor cells (OPCs) have high oxidative phosphorylation (OXPHOS) rates before and during myelination, and myelinating oligodendrocytes have low OXPHOS and high glycolytic rates^1,2^. The mechanisms regulating this metabolic switch are not well understood. However, mitochondria undergo morphological remodeling during maturation *in vivo*, supporting a shift in metabolic demands^3^. The dependence on fatty acid oxidation (FAO) during this maturation stage is not well understood in oligodendrocytes. Mitochondrial FAO generates acetyl-CoA to initiate the tricarboxylic acid (TCA) cycle. Cells typically depend on FAO when glucose levels in the cell are low. Previous studies have shown that FAO is required for neural stem cell maintenance and proper neurogenesis^4^. Fatty acid β-oxidation in oligodendrocytes supports axonal energy metabolism when glucose is depleted, indicating that oligodendrocyte fatty acid metabolism can serve as an energy reserve for the bioenergetic needs of these cells^5^.

Myeloid cell leukemia-1 (MCL-1) is a well characterized anti-apoptotic protein, known to inhibit mitochondrial-mediated cell death. MCL-1 has non-apoptotic functions which are critical for post-embryonic development^6^. We have previously shown that MCL-1 regulates mitochondrial morphology and dynamics of human pluripotent stem cells as well as other differentiated cells such as cardiomyocytes^7,8^. This non-apoptotic function of MCL-1 is critical for maintaining the bioenergetic needs of cells during differentiation. MCL-1 also promotes long-chain fatty acid β-oxidation by interacting with long-chain acyl-CoA synthetase 1 (ACSL1), an enzyme that converts long-chain fatty acids into acyl-CoAs^9^. This interaction is disrupted by the MCL-1 inhibitor and BH-3 mimetic, S63845^9,10^. We have previously shown that knocking out *MCL-1* in mouse neural precursor cells leads to a decrease in myelinating oligodendrocytes^11^. Together, these findings suggest a potential link between oligodendrocyte function and the non-apoptotic role of MCL-1.

To investigate the mechanisms that modulate the development of human oligodendrocytes, we generated and optimized an *in vitro* system to isolate and co-culture human embryonic stem cell (hESC)-derived OPCs that undergo maturation when co-cultured with motor neurons^12^. This system allows for the investigation of downstream effects on mitochondrial morphology, gene expression of FAO enzymes, and expression of oligodendrocyte-associated genes following treatment. Additionally, using the co-culture system, we found that pharmacological inhibition of MCL-1 at the OPC stage leads to morphological changes in oligodendrocytes expressing myelin basic protein (MBP) and alterations to their mitochondria morphology. This is the first report demonstrating remodeling of mitochondrial morphology during the development of human oligodendrocytes.

## Results

### Generation of the oligodendrocyte lineage from human stem cells

We optimized a protocol for the generation of myelinating oligodendrocytes from stem cells that can be used to manipulate and investigate oligodendrocyte lineage cells at the various stages of development. A mixed culture consisting of OPCs, neurons, and astrocytes was generated using a series of small molecule and growth-factor-defined media^13^. To purify the population of OPCs, we used the MACS cell separation system with Anti-A2B5 MicroBeads at 65 days *in vitro* (DIV) (**Figure 1A**). OLIG2-positive cells were on average 72% of the cell population following the cell sort (**Supplementary Figures 1A and 1B**). To further elucidate the non-apoptotic function of MCL-1 during the development of oligodendrocytes, we treated isolated OPCs with S63845 for 48-hours. Previous studies outlining the mechanism of action of S63845 found that inhibition of MCL-1 leads to increased protein stability, consequently increasing protein levels^10^. Following the treatment of OPCs, protein levels of MCL-1 increased as anticipated, confirming the effectiveness of pharmacological MCL-1 inhibition (**Figure 1B**).

**Figure 1.**
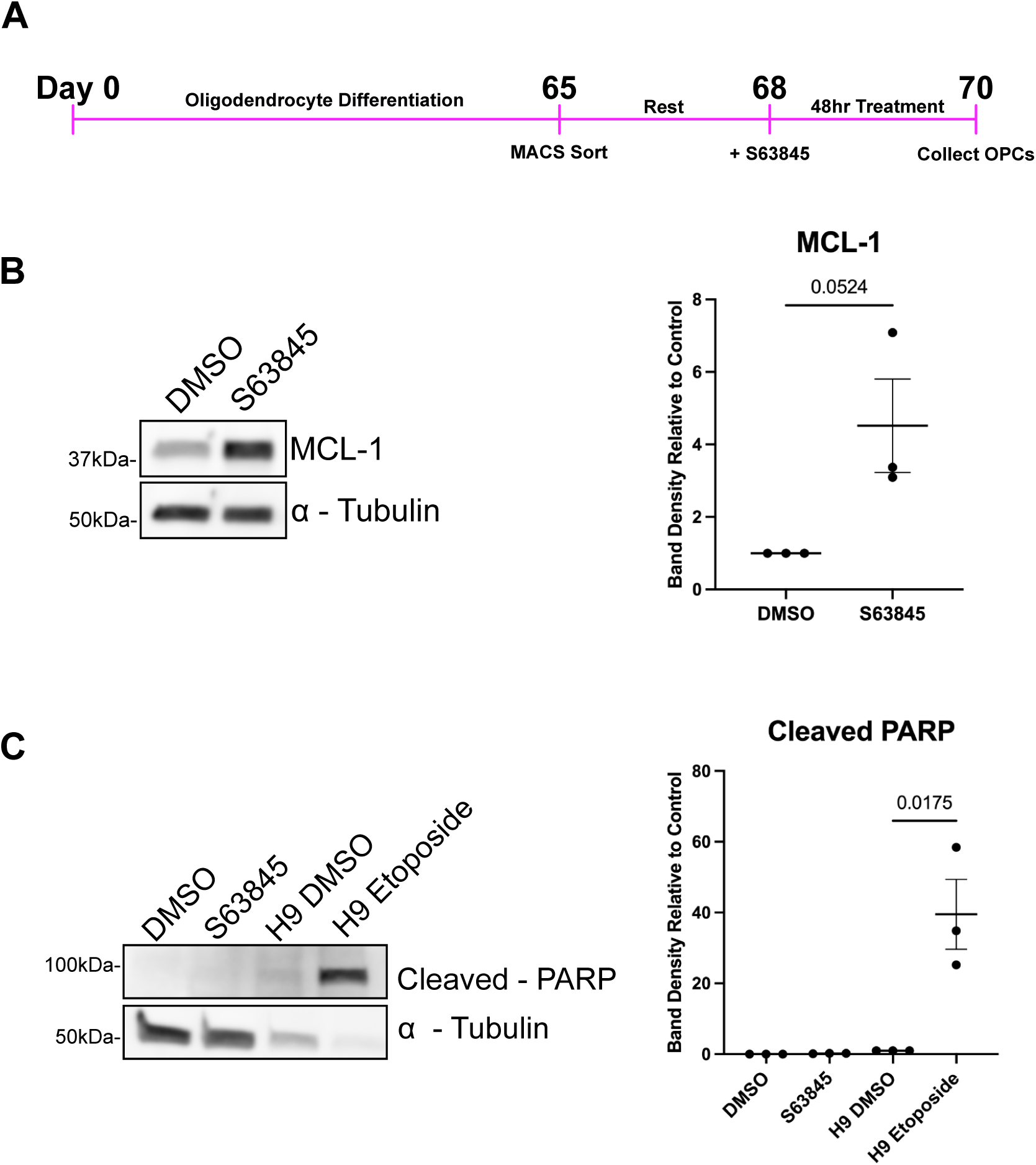
*In vitro* human oligodendrocyte system reveals the non-apoptotic effect of MCL-1 inhibition. (**A**) Timeline of oligodendrocyte differentiation and treatment with S63845. At day 65 of differentiation, cells undergo cell sorting using the MACS cell separation system. Following a rest period, cells were treated with DMSO (vehicle) and S63845 for 48 hrs. At the 48-hour timepoint, cells were collected. (**B**) Western blot of MCL-1 levels in OPCs following DMSO (vehicle) and S63845 treatment (left panel) and band density quantification (right panel). Each dot on graphs represents a biological replicate (n=3), analyzed by Student’s t-test, and error bars represent mean ± SEM. (**C**) Western blot of cleaved PARP following S63845 treatment in OPCs and etoposide treatment in H9 human embryonic stem cells, respectively (left panel) and band density quantification (right panel). Both treatments used DMSO as vehicle control. Each dot on graphs represents a biological replicate (n=3), analyzed by Student’s t-test, error bars represent mean ± SEM.

### MCL-1 inhibition does not lead to cell death in oligodendrocyte precursor cells

MCL-1 is a highly regulated protein that inhibits cell death by interacting with pro-apoptotic proteins through its BH3 domain^14,15^. The anti-apoptotic function of MCL-1 has a specific temporal requirement during early neurogenesis, and it has been shown that it is necessary for the survival of neural progenitor cells in the developing mouse brain^16,17^. However, the role of MCL-1 during oligodendrogenesis is not known. Recent findings show that the apoptotic function of MCL-1 is critical for embryonic development, and the non-apoptotic function of MCL-1 is needed for post-embryonic development^6^. Our findings here demonstrate that MCL-1 is dispensable for the survival of OPCs. Following treatment of OPCs with S63845 for a 48-hour period, protein levels of cleaved-PARP, an indicator of caspase activation, were not altered (**Figure 1C**). As a positive control to validate the protein levels of cleaved-PARP, we treated (hESCs) (H9 line) with etoposide, a DNA-damaging reagent that leads to cell death. Following treatment with etoposide, cleaved-PARP levels were increased, indicating caspase-mediated apoptosis. The absence of cell death following pharmacological inhibition of MCL-1 indicates that the findings reported here are independent of a cell death phenotype.

### MCL-1 inhibition does not alter mitochondrial morphology in oligodendrocyte precursor cells

Mitochondria are dynamic organelles constantly undergoing coordinated fusion and fission events needed to maintain mitochondrial function^18^. Mitochondrial fission events are largely mediated by dynamin-related protein 1 (DRP1), while fusion events are mediated by mitofusin 1/2 (MFN1/2) and optic atrophy 1 (OPA1)^19^. We and others have shown that alterations to DRP1 can lead to detrimental changes in mitochondrial function and, subsequently, neuronal and glial function^20–22^. Thus, it is critical to understand the tight regulation of these processes during neuronal and glial development as they have broader contributions to cellular function. The modifications of mitochondrial dynamics and ultrastructure determine the metabolic state of the cell and its adaptation to a new state^23–25^.

We have previously, shown that inhibition of MCL-1 leads to mitochondrial elongation in hESCs and fragmentation in cardiomyocytes derived from human induced pluripotent stem cells through the potential interplay with DRP1 and OPA1^7,8^. These findings indicate that MCL-1 helps to maintain mitochondrial morphology, which is essential for energy homeostasis. The role of MCL-1 on mitochondrial function during oligodendrocyte development is largely unknown. Here, we treated isolated OPCs with S63845 for 48- hours and did not see significant morphological changes to the mitochondria. We used oligodendrocyte transcription factor 2 (OLIG2) and platelet derived growth factor receptor alpha (PDGFRα) as markers to identify OPCs following treatment (**Figure 2A**)^26,27^. This co-staining method allowed for visualization of the mitochondrial network within the entire cell. There were no significant changes to the mitochondrial surface area, volume, sphericity, diameter, or major axis length (**Figures 2B-2F**).

**Figure 2.**
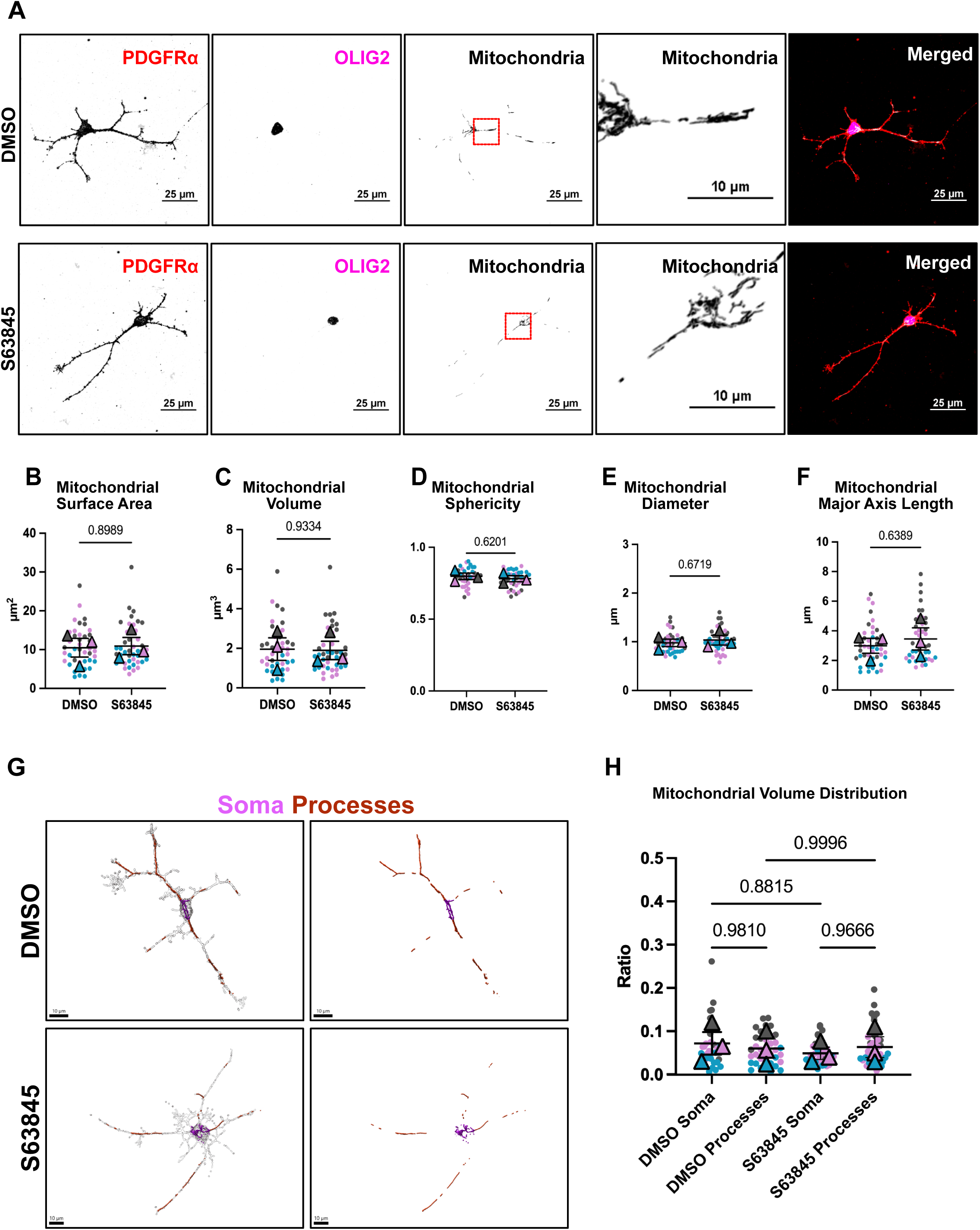
Pharmacological inhibition of MCL-1 does not cause alterations to mitochondrial morphology in OPCs. (**A**) Representative spinning disk confocal maximum intensity projections of immunofluorescent staining for platelet-derived growth factor receptor alpha (PDGFRα) (red), oligodendrocyte transcription factor 2 (OLIG2) (magenta), and mitochondria (white) in DMSO (vehicle) and S63845 treated OPCs (scale bar = 25µm). Zoom-in of perinuclear mitochondria (scale bar = 25µm). (**B-F**) Quantification of mitochondrial surface area, volume, sphericity, diameter, and major axis length. Each color represents a biological replicate (n=3), each dot represents a cell (10-15 per n), each triangle represents the mean of the biological replicate, analyzed by Student’s t-test, and error bars represent mean ± SEM. (**G**) 3D reconstructions of OPCs and their mitochondrial network treated with DMSO (vehicle) and S63845. Mitochondria in magenta are in the cell’s soma and mitochondria in red are in the cell’s processes. (**H**) Distribution of mitochondria in OPCs. Ratio calculated by diving the volume of the mitochondria in the soma (shown in magenta in Figure 2G) or processes (shown in red in Figure 2G) by the total volume of the OPC. Each color represents a biological replicate (n=3), each dot represents a cell (10-15 per n), each triangle represents mean of biological replicate, analyzed by one-way ANOVA, error bars represent mean ± SEM. Conditions were blinded to experimenter for 3D reconstructions.

This is the first report, to our knowledge, of mitochondrial morphology remodeling in human OPCs in an *in vitro* system. The mitochondria in OPCs appear elongated, and its network expands throughout the soma and processes of the cell. The mitochondria appear to wrap around the nucleus and continue to reach into the processes where they may be needed to meet the energetic demands of the growing cell. Bame and Hill reported an *in vivo* mouse system to describe mitochondrial network reorganization throughout the development of oligodendrocytes^3^. The characterization of the mitochondrial morphology recapitulates their findings in the OPC stage. To compare the two model systems, we used a similar analytic workflow that allows for comparison of the mitochondrial volume in the soma and processes of the OPC. After generating 3D reconstructions of OPCs using Imaris image analysis software, we developed a ratio to quantify the volume of mitochondria within the soma or processes normalized to the total volume of the OPC (**Figure 2G**). We found no significant differences in the mitochondrial distribution between the soma and the processes (**Figure 2H**). Inhibition of MCL-1 did not modify the distribution of mitochondria within the cell. Additionally, there are no significant differences in the volume of PDGFRα positive cells (**Supplementary Figure 2**). Consequently, we can speculate that pharmacological inhibition of MCL-1 does not affect the distribution of mitochondria within OPCs.

### MCL-1 inhibition in human oligodendrocyte precursor cells may lead to alterations in the fatty acid β-oxidation pathway

OPCs are critical during oligodendrogenesis as they proliferate and simultaneously integrate into the developing neuronal system^27^. At this stage, they have high energetic demands as they extend their processes to search for axons and prepare to generate myelin sheaths, which are lipid-rich structures^29^. OPCs rely on OXPHOS and switch to glycolysis as they mature into myelinating oligodendrocytes^1,2^. Previous findings established that oligodendrocytes depend on fatty acid β-oxidation in low glucose conditions^5^. Thus, ATP produced from the oxidation of fatty acids in the mitochondria is sufficient to maintain the function of the oligodendrocyte when glycolysis is not available. However, the role of FAO during this developmental switch is not well understood. MCL- 1 regulates long-chain fatty acid β-oxidation by its interaction with ACSL1^9^. Activation of fatty acids by ACSL1 triggers a cascade of events regulated by CPT1 and CPT2, two key enzymes that work together to transport fatty acids into the mitochondria^30^. Once inside the mitochondria, fatty acyl-CoA undergoes β-oxidation to generate acetyl-CoA for the TCA cycle^31–33^ (**Figure 3A**). It has been previously reported that FAO genes are downregulated following genetic deletion of *Mcl-1* in mouse B-cell lymphoblastic leukemia cells and liver tissue^34^. To understand if this regulatory mechanism is conserved in OPCs, we examined changes in gene expression of *MCL1*, *ACSL1*, *CPT1*, and *CPT2* following inhibition of MCL-1. Gene expression of *MCL1*, *CPT1A*, and *CPT2* was not altered; however, *ACSL1* was significantly decreased (**Figure 3B-3E**). These results indicate that MCL-1 may regulate fatty acid metabolism in OPCs as alterations to *ACSL1* expression could have downstream effects on the activation of fatty acids.

**Figure 3.**
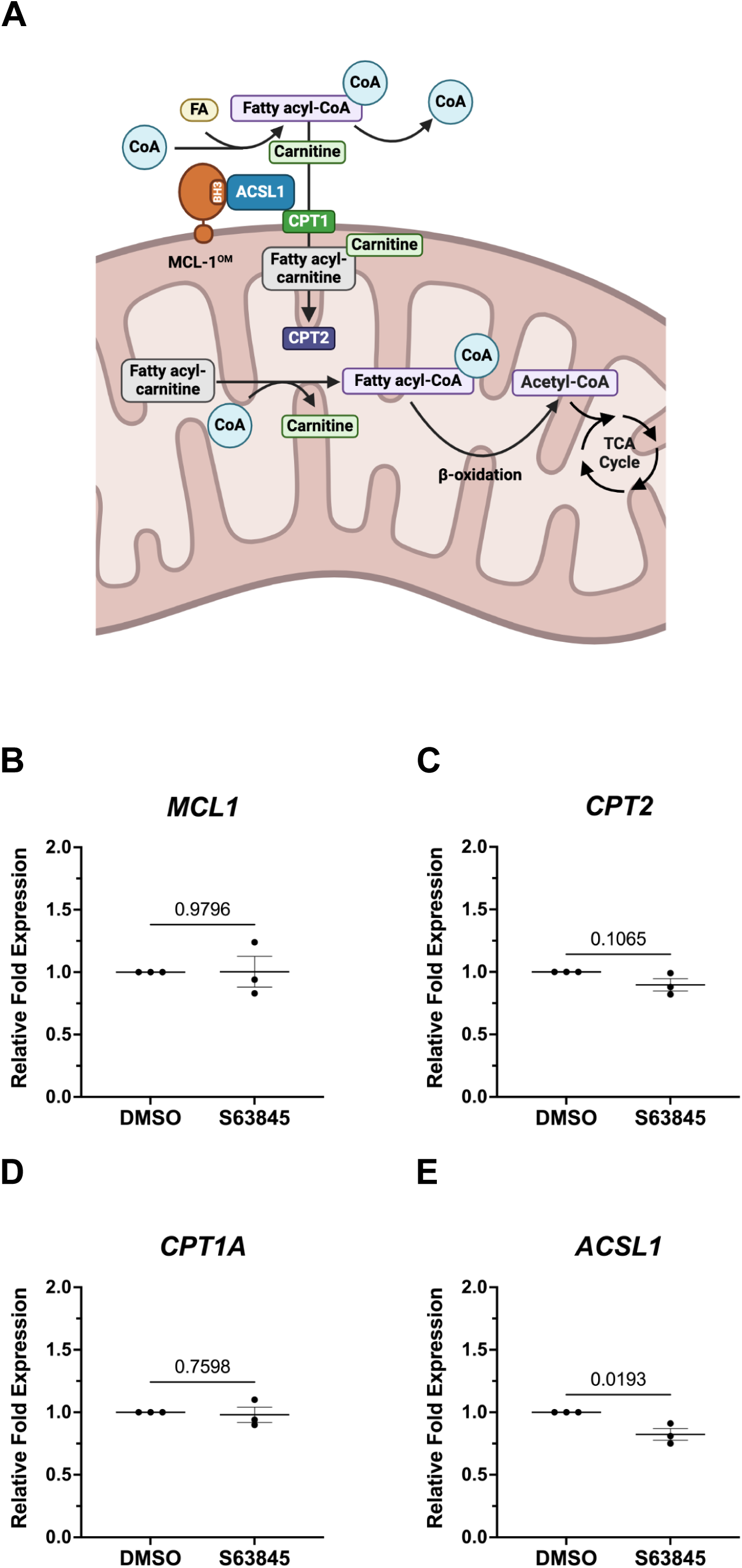
Pharmacological inhibition of MCL-1 leads to decreased gene expression of *ACSL1*. (**A**) Schematic outlining activation of fatty acids (FA) and subsequent transport into the mitochondrial matrix for oxidation. MCL-1 at the outer mitochondrial membrane (MCL- 1^OM^) may associate with ACSL1promoting its function. ACSL1 catalyzes the conversion of long-chain fatty acids to their activated form, fatty acyl-CoA, which can cross the outer mitochondrial membrane and enter the intermembrane space. To be transported into the mitochondrial matrix, fatty acyl-CoA must be converted to fatty acyl-carnitine. Carnitine palmitoyltransferase 1 (CPT1) catalyzes the transfer of the acyl group of a fatty acyl-CoA from coenzyme A to carnitine, allowing fatty acyl-carnitine to enter the matrix. Once inside the mitochondria matrix, carnitine palmitoyltransferase 2 (CPT2) converts fatty acyl-carnitine back to fatty acyl-CoA. β-oxidation breaks down fatty acyl-CoA to acetyl-CoA. Acetyl-CoA molecules then enter the tricarboxylic acid (TCA) cycle to generate ATP. (**B- E**) qPCR relative fold expression of *MCL1*, *ACSL*, *CPT1A*, and *CPT2*. Each dot on graphs represents a biological replicate (n=3) consisting of the average of two technical replicates, analyzed by Student’s t-test; error bars represent mean ± SEM.

### Inhibition of MCL-1 in human oligodendrocyte precursor cells leads to decreased expression of key oligodendrocyte-related genes

SRY-box transcription factor 10 (SOX10) and OLIG2 are both key transcription factors that are involved in the development and maturation of oligodendrocytes^35–37^. OLIG2 is expressed early in ventral progenitor cells, and SOX10 is present in the pre-OPC to myelinating oligodendrocyte stages^38,39^. SOX10 and OLIG2 modulate each other to control oligodendrocyte differentiation^40–42^. Expression of both OLIG2 and SOX10 at the OPC stage ensures that transcriptional regulatory networks are functional to maintain oligodendrocyte lineage identity^38^. MBP is the most abundant protein in compact myelin^43^. It is responsible for interacting with lipids to maintain the structure of the myelin sheath^44–46^. Loss of MBP leads to severe dysmyelination in the central nervous system^47,48^.

Previous findings from our group have revealed that conditional deletion of *Mcl-1* in neural progenitor cells leads to loss of SOX10 and MBP in the developing mouse brain^11^. We measured gene expression of *SOX10*, *OLIG2*, *PDGFRA*, and *MBP* following pharmacological inhibition of MCL-1 in OPCs. Gene expression of *OLIG2* and *MBP* were significantly downregulated; however, expression of *SOX10* and *PDGFRA* were maintained (**Figure 4A-D**). There were no changes in protein levels of SOX10 and OLIG2 following treatment (**Supplementary Figure 3A-B**). Loss of OLIG2 can arrest differentiation, leading to changes in the capacity of OPCs to mature into myelinating oligodendrocytes and establish connections with axons to generate compact sheaths^49,50^. We speculated that alterations in gene expression of *OLIG2* and *MBP* at the OPC-state may contribute to the function and maturation of oligodendrocytes later in development. Moreover, possible changes to metabolic support following inhibition of MCL-1 may lead to alterations in the ability of OPCs to transition into myelinating oligodendrocytes.

**Figure 4.**
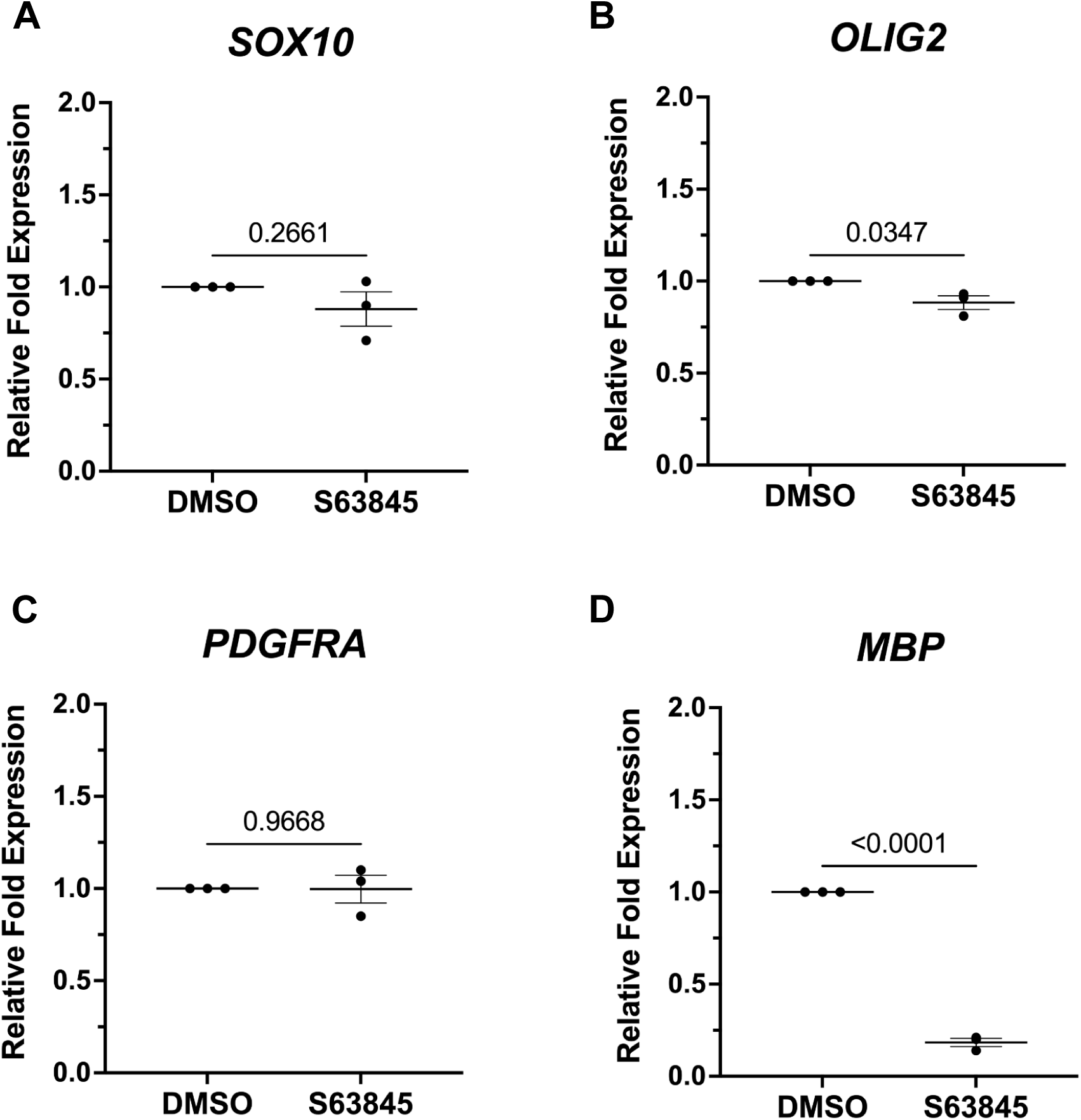
Pharmacological inhibition of MCL-1 leads to decreased gene expression of *OLIG2* and *MBP*. (**A-D**) qPCR relative fold expression of *SOX10*, *OLIG2*, *PDGFRA*, and *MBP*. Each dot on graphs represents a biological replicate (n=3) consisting of the average of two technical replicates, analyzed by Student’s t-test, error bars represent mean ± SEM.

### Inhibition of MCL-1 at early stages of differentiation alters mitochondrial morphology in oligodendrocytes

It has been shown that OPCs reorganize their mitochondrial network as they mature into myelinating oligodendrocytes *in vivo*^3^. In mice, myelinating oligodendrocytes have decreased mitochondrial volume compared to OPCs, and the mitochondria shift to a fragmented state^3,25^. This dramatic reorganization may be attributed to the shift in metabolic state that occurs with the growth of the cytoplasmic content and the formation of myelin sheath. To examine the mitochondrial morphology in mature oligodendrocytes, we used a co-culture system^12^. Immediately following a 48-hour treatment with S63845, OPCs were cultured with human induced pluripotent stem cell-derived motor neurons (**Supplementary Figure 4A-B**). The co-culture was maintained for 20 DIV and then fixed for imaging analysis (**Figure 5A**). To capture the entire cell, the co-culture was stained with myelin oligodendrocyte glycoprotein (MOG), which is found on the cell surface of myelinating oligodendrocytes and external lamellae of myelin sheaths^51^. MOG, along with SOX10 staining, allowed for the identification of mature cells in the cell culture system and the generation of 3D reconstructions. As shown in the murine model, human oligodendrocytes have fragmented mitochondria (**Figure 5B**)^3,25^. There are no significant changes to the mitochondrial surface area, volume, sphericity, and diameter (**Figures 5C-F**) in oligodendrocytes previously treated with S63845. However, pharmacological inhibition of MCL-1 at the OPC stages leads to significantly decreased mitochondrial major axis length in mature oligodendrocytes (**Figure 5G**). This alteration to mitochondrial morphology in oligodendrocytes suggests that MCL-1 is required to regulate the metabolic state of the cell during differentiation. To assess mitochondrial distribution in mature oligodendrocytes, we measured the volume of mitochondria within the soma or processes and divided it over the total volume of the oligodendrocyte to generate a ratio. Results indicate no significant alterations in the mitochondrial distribution between the soma and the processes in vehicle-treated cells (**Figure 5H-I**) and no significant differences in the volume of MOG-positive cells (**Supplementary Figure 5**).

**Figure 5.**
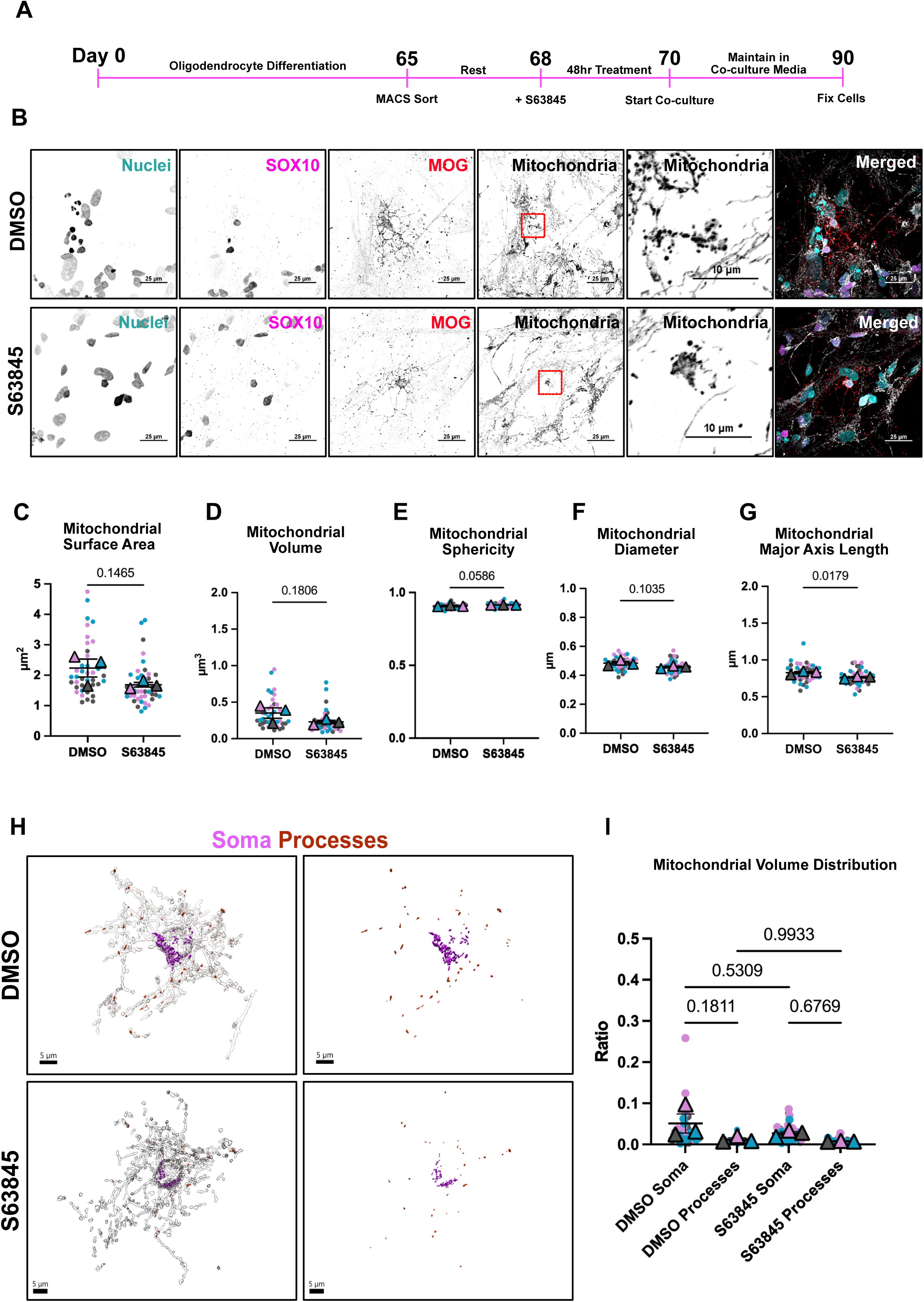
Pharmacological inhibition of MCL-1 at early stages of development leads to mitochondrial morphology alterations in mature oligodendrocytes. (**A**) Timeline of oligodendrocyte differentiation, treatment with S63845, and co-culture with motor neurons. At day 65 of differentiation, cells undergo cell sorting using the MACS cell separation system. Following a rest period, cells were treated with DMSO (vehicle) and S63845 for 48hrs. At the 48hr timepoint, cells were cultured with motor neurons. Co- culture was maintained until day 90. (**B**) Representative spinning disk confocal maximum intensity projections of immunofluorescent staining for nuclei (cyan), SOX10 (magenta), myelin oligodendrocyte glycoprotein (MOG) (red), and mitochondria (white) in mature oligodendrocytes that were previously treated with DMSO (vehicle) and S63845 (scale bar = 25µm). Zoom in of mitochondria at the nucleus (scale bar = 25µm). (**C-G**) Quantification of mitochondrial surface area, volume, sphericity, diameter, and major axis length. Each color represents a biological replicate (n=3), each dot represents a cell (10- 15 per n), each triangle represents mean of biological replicate, analyzed by Student’s t- test, error bars represent mean ± SEM. (**H**) 3D reconstructions of mature oligodendrocytes and their mitochondrial network treated with DMSO (vehicle) and S63845. Mitochondria in magenta are in the cell’s soma and mitochondria in red are in the cell’s processes. (**I**) Distribution of mitochondria in mature oligodendrocytes. Ratio calculated by dividing the volume of the mitochondria in the soma (shown in magenta in Figure 6G) or processes (shown in red in Figure 6G) by the total volume of the mature oligodendrocyte. Each color represents a biological replicate (n=3), each dot represents a cell (10-15 per n), each triangle represents mean of biological replicate, analyzed by one-way ANOVA, error bars represent mean ± SEM. Conditions were blinded to the experimenter for 3D reconstructions

### Inhibition of MCL-1 at early differentiation stages alters the morphogenesis of oligodendrocytes cells without disrupting axon contacts

To examine if changes to gene expression of *OLIG2* and *MBP* had an effect on the capacity of the OPCs to mature, we used the motor neuron co-culture system described previously^12^. Axons from motor neurons were stained for neurofilament 200 (NF-200), and oligodendrocytes were stained for MBP. 3D reconstructions were generated with Imaris, allowing for detailed visualization of oligodendrocytes and their processes. We used the Imaris XTension, Surface-Surface Contact Area, to visualize and quantify the contact between oligodendrocytes and axons (**Figure 6A**). There is a significant decrease in the volume of oligodendrocytes previously treated with S63845 and a downward trend in the cell’s area (Figure 6B-C). We generated a ratio of the volume or area of the oligodendrocyte divided by the volume or area of the surface contact between the oligodendrocyte and surrounding axons (termed oligodendrocyte–axon contact). The oligodendrocyte–axon contact volume and area are similar between the vehicle and S63845 treated groups (**Figure 6D-E**). Sholl analysis of MBP+ cells revealed that oligodendrocytes in the S63845 treated group have significantly fewer intersections at 30µm and 40µm distance from cell center, suggesting a loss in morphological complexity of these cells (**Figure 6F-G**). Th loss of volume coupled with decreased complexity suggests that the non-apoptotic function of MCL-1 at the OPC stage is required for the morphogenesis of oligodendrocytes. Possible alterations to FAO and the subsequent ATP production may not be sufficient to support the bioenergetic demands required during oligodendrocyte maturation. At the early stage of myelination examined here, oligodendrocyte processes have the capacity to contact axons. The alterations in morphogenesis may contribute to the ability of the cell to adequately make axonal contacts later in development.

**Figure 6:**
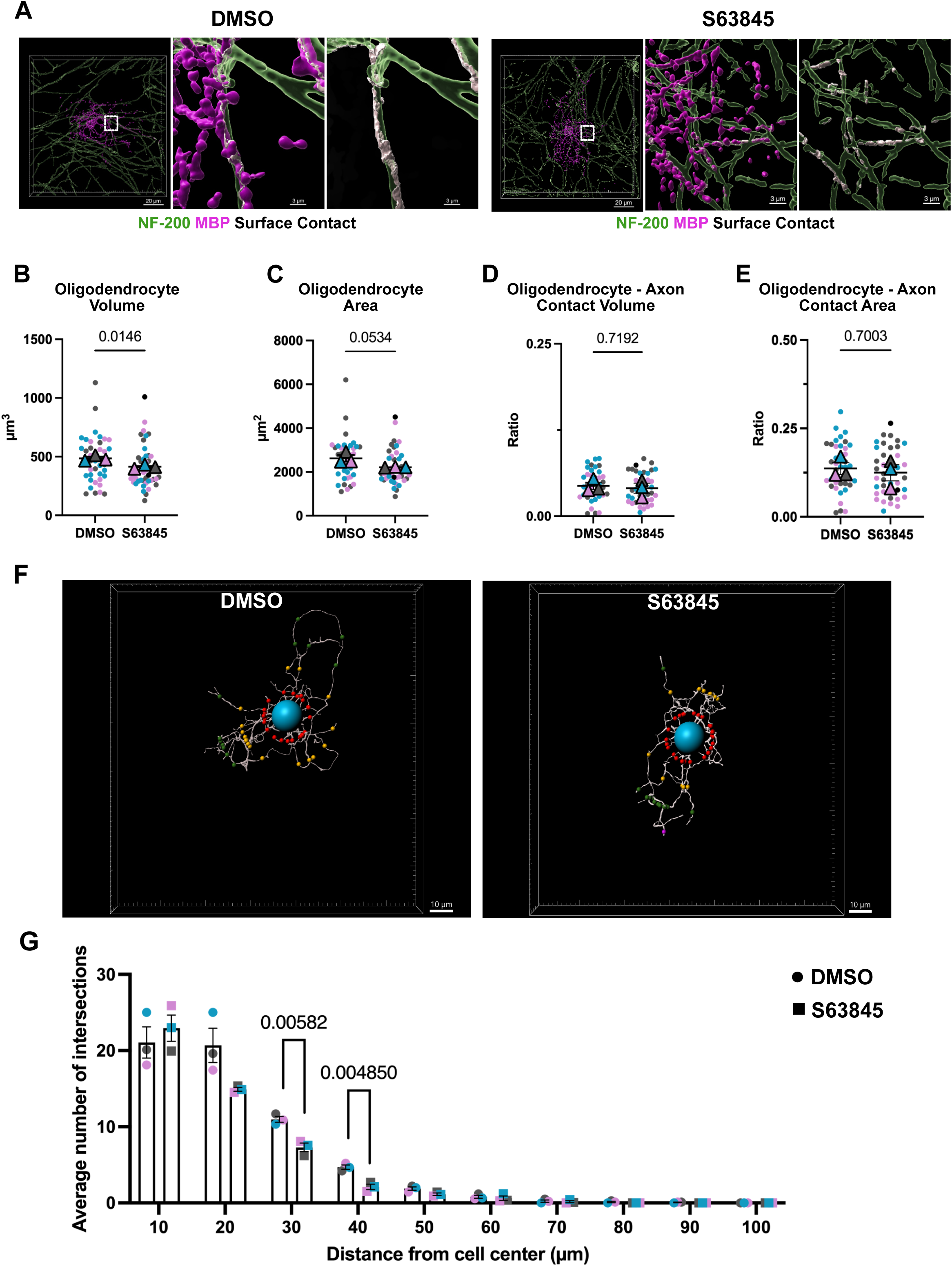
Using a human *in vitro* co-culture system to investigate the long-term effects of pharmacological inhibition of MCL-1 in OPCs. (**A**) 3D reconstructions of myelin basic protein (MBP) positive cells (magenta), surrounding axons (green), and surface contact (white) in DMSO (vehicle) and S63845 treated cells. Scale bar = 20µm. Scale bar = 3µm in zoomed-in panels. (**B-C**) MBP positive oligodendrocyte volume and area in DMSO (vehicle) and S63845 treated cells. Each color represents a biological replicate (n=3), each dot represents a cell (10-15 per n), each triangle represents the mean of biological replicate, analyzed by Student’s t-test, error bars represent mean ± SEM. Conditions were blinded to the experimenter for 3D reconstructions. (**D**) Ratio calculated by dividing the surface contact volume (white in Figure 5B) by the oligodendrocyte volume (magenta in Figure 5B). Each color represents a biological replicate (n=3), each dot represents a cell (10-15 per n), each triangle represents the mean of the biological replicate, analyzed by Student’s t-test, error bars represent mean ± SEM. Conditions were blinded to the experimenter for 3D reconstructions. (**E**) Ratio calculated by dividing the surface contact area (white in Figure 5B) by the oligodendrocyte area (magenta in Figure 5B). Each color represents a biological replicate (n=3), each dot represents a cell (10-15 per n), each triangle represents the mean of the biological replicate, analyzed by Student’s t-test, error bars represent mean ± SEM. Conditions were blinded to the experimenter for 3D reconstructions. (**F**) 3D filament reconstruction of MBP positive oligodendrocytes. Sholl analysis is indicated by different colored dots placed every 10µm from cell center (blue dot). Conditions were blinded to the experimenter for 3D filament reconstructions. (**G**) Sholl analysis histogram demonstrating the average number of intersections (y-axis) and distance from the cell’s center (x-axis). Each dot or square represents the mean number of intersections at a given distance (mean of 10-15 cells per biological replicate) in DMSO (vehicle) or S63845 treated cells, respectively. Each color represents a biological replicate (n=3). Analyzed by Student’s t-test, error bars represent mean ± SEM.

## Discussion

The developing brain has a high energic demand that relies on glucose and lactate for normal function^52,53^. Although understudied, it is known that the oxidation of fatty acids is largely responsible for the brain’s energy consumption as well^54,55^. Loss of lipid homeostasis has been described in Alzheimer’s Disease pathology, illustrating the importance of this mechanism for human health^56^. The balance and interactions between glycolysis and FAO in the developing brain are poorly understood. FAO of short, medium, and long-chain fatty acids occurs in the mitochondria^57,58^. MCL-1 is thought to modulate long-chain fatty acid β-oxidation by its interaction with ACSL1^9^. However, it is not known if this non-apoptotic function of MCL-1 is conserved in other cell types, such as oligodendrocytes, which are known to depend on MCL-1 for proper development^11^. Mature oligodendrocytes are mainly glycolytic yet require FAO to uphold the metabolic support provided to axons^1,2,5^. Here, we describe a platform to investigate human oligodendrocyte development. This platform allows for the isolation of OPCs to examine their potential to maturate into myelinating oligodendrocytes. Outlining cell transition states in early development is critical for understanding key elements of differentiation and integration of oligodendrocytes into the neuronal system.

Proper integration of oligodendrocytes into the developing brain requires high energy production to support the expanding lipid-rich membrane that composes the myelin sheath^53^. Alterations to this process may lead to hyper- or hypomyelination, triggering severe white matter disorders^59,60^. Regulation of mitochondrial dynamics and morphology is critical in supporting the cellular and energetic demands of oligodendrocytes during development^25^. Here, we present the first report of the characterization of mitochondrial morphology changes in an *in vitro* human system of oligodendrogenesis. Similar to findings in mouse tissue, mitochondria in OPCs have an elongated morphology and fragment as the cells mature^3,25^. This is accompanied by an apparent loss of mitochondrial volume in mature oligodendrocytes. These findings fit the concept that oligodendrocytes switch to glycolysis from OXPHOS, as glycolytic cells are thought to have fragmented mitochondria and cells that rely on OXPHOS have elongated mitochondria^61^. However, the association between mitochondrial morphology and metabolic state is dependent on the ultrastructure of the mitochondria, since mitochondria with an increased volume of cristae have a higher capacity for efficient ATP production^62^.

As OPCs develop and extend processes to explore the surrounding environment in search of axons to myelinate, it is expected to see a shift in mitochondrial distribution^63^. However, the absence of neurons in the OPC culture post-sorting may prevent the growth of the processes, leading them to not have the same response as previously described *in vivo*^29^. Additionally, pharmacological inhibition of MCL-1 did not alter mitochondrial morphology or distribution in OPCs, suggesting that MCL-1 inhibition does not affect the mitochondrial morphology at this stage of development. However, ultrastructural alterations to the mitochondria, which are further indicative of the energetic state of the cell, require further examination^64^.

We found that inhibition of MCL-1 leads to decreased gene expression of oligodendrocyte-related genes, *OLIG2* and *MBP*. Acetyl-CoA derived from FAO is a major contributor to the acetyl-CoA pool^65^. Maintaining the acetyl-CoA pool is vital for TCA cycle activity and the metabolites that regulate chromatin modifications and DNA methylation^66^. Since MCL-1 controls long-chain fatty acid β-oxidation, we speculate that inhibition of MCL-1 may lead to a decrease in the acetyl-CoA pool and subsequent modifications to gene regulation that can impact oligodendrocyte fate, function, and maturation. We also detected a dramatic loss in the volume of MBP positive cells, but not MOG positive cells. These data demonstrate a possible dysfunction in myelin integrity as MBP is responsible for interacting with lipids to form tight adhesions of the cytosolic surfaces^44^. The resolution provided by the spinning disk confocal images and 3D reconstructions presented here reveals that oligodendrocytes may still be in the early stages of development. Thus, the long-term maintenance of the myelin sheath and axonal contact should be further investigated. Additionally, it would be informative to investigate if changes to FAO leads to alterations in established oligodendrocyte–axon contacts, as the energic requirements in myelinating oligodendrocytes differ from OPCs.

In our system and previously reported systems, it is technically challenging to isolate a pure population of human OPCs and maintain them for a long period of time. This limitation has led to constraints on the experiments that can be performed to further investigate FAO, such as respirometry assays, metabolomics, and carbon tracing. Improved techniques are needed in the field of glia biology to further elucidate metabolic requirements that regulate the development of human oligodendrocytes as they can best capture disease-relevant features^67,68^. Diseases like multiple sclerosis (MS), leukodystrophies, and other demyelinating disorders affect human oligodendrocytes specifically. While murine models are invaluable for understanding basic cellular processes, they often fail to fully capture the complexity of these diseases due to species differences in immune responses, neural development, and cell interactions. Isolating and maintaining human oligodendrocytes with myelination capacity *in vitro* facilitates the investigation of human-specific pathophysiology, drug responses, and disease mechanisms.

The striking parallels in mitochondrial network reorganization between human and mouse models suggest that the core mechanisms governing mitochondrial morphology and function during the development of oligodendrocytes may be evolutionarily conserved. Additionally, these results imply that specific mitochondrial events, such as the mitochondrial transport along oligodendrocyte projections vital for sustaining long-distance signaling and cellular metabolism, appear to operate similarly in both species. Thus, integrating both models provides a more holistic approach to studying mitochondrial function and enables the identification of conserved mitochondrial mechanisms regulating oligodendrogenesis. The detection of species-specific differences could also inform the development of more targeted and effective therapies for mitochondrial-related disorders in humans.

## Author contributions

M. Gil and V. Gama conceived the study, designed experiments, interpreted data, and wrote the manuscript. V. Gama supervised and funded the research. M. Gil designed and carried out all the cell biology experiments with technical assistance from M. Hanna. All authors edited the document.

## Acknowledgments

This work was supported by 2R35GM128915-06 (VG) and HHMI Gilliam Fellowship GT15720 (MG). Data analysis and figure creation were partly performed using the Vanderbilt Cell Imaging Shared Resource (supported by NIH grants CA68485, DK58404, and EY08126). We thank Madison Yarbrough for assisting with blinding conditions for image analyses and David Carmona Berrio for providing technical guidance. Figure 3A in this manuscript was created using BioRender (BioRender.com). The authors declare no competing financial interests.

## Methods

### Cell line and stem cell maintenance

H9 human embryonic stem cells (WiCell Research Institute, WA09, NIH Registration Number: 0062) were maintained in mTeSR1 (STEMCELL Technologies, 85850) on Matrigel-coated (Corning, 354277) plates. Media was changed daily and passaged when they reached when they reached 60-80% confluency.

### Etoposide Treatment

H9 hESCs were treated with 20uM etoposide (Sigma-Aldrich, E1383) for 24hrs. DMSO was used as vehicle control.

### Oligodendrocyte differentiation

Differentiation was adapted from Douvaras and Fossati, 2015^13^. Detailed instructions for the differentiation can be found at Gil and Gama 2024^12^. Basal media was used throughout the differentiation and composed of DMEM/F12 (Thermo Fisher Scientific, 11320033), 1X Penicillin-Streptomycin (Thermo Fisher Scientific, 15-140-122), 1X MEM Non-Essential Amino Acid (Thermo Fisher Scientific, 11-140-050), 1X GlutaMAX (Thermo Fisher Scientific,), and 55 µM 2-Mercaptoethanol (Bio-Rad, 1610710).

H9 hESCs were dissociated using Accutase (Thermo Fisher Scientific, A1110501) and incubated at 37 °C for 5 min. Accutase was diluted with DMEM/F12 medium. A cell lifter was used to remove cells from the well and fully dissociate by pipetting mixture 2–5 times with a p1000 pipette. Cells were centrifuged at 200× g for 4 min at RT. Cells were plated at 8 × 10^4^ cells per well on a Matrigel-coated 6-well plate in mTeSR1 supplemented 10 µM Y27632 (STEMCELL Technologies, 72307). The following day, mTeSR1 was replenished to remove Y27632. Medium changes were performed daily until 2 days following replating when cells reached 80% confluency. At this point, differentiation was started (Day 0) by feeding cells with neural induction medium (NIM) composed of basal media supplemented with 25 µg/mL insulin (Sigma-Aldrich, I9278), 10 µM SB431542 (Reprocell, 04-0010-10), 250 nM LDN193189 (Reprocell, 04-0074), and 100 nM Retinoic acid (Sigma-Aldrich, R2625). Media was changed daily and SB431542, LDN193189, and Retinoic acid were added fresh each day. On Day 8, cells were fed with N2 medium. N2 medium was composed of basal medium with 1X N2 supplement (Thermo Fisher Scientific, 17502048), and 250 nM SAG Hedgehog pathway activator (STEMCELL Technologies, 73412). Medium was changed daily until Day 12. Retinoic acid and SAG were added fresh each day. On Day 12, aggregates were formed by mechanical dissociation. Old medium was removed from wells and replenished with 1mL of N2B27 medium. N2B27 medium was composed of basal media supplemented with 1X N2 supplement, 1X B27 minus vitamin A supplement (Thermo Fisher Scientific, 12587010), 25 µg/mL insulin, 100 nM Retinoic acid, and 250 nM SAG. Aggregates were generated by using a cell lifter placed perpendicular to the bottom of the well and creating 20 parallel cuts into the cell layer to cover the entire well. Plate was turned 90° and 20 more cuts were made. Finally, plate was turned 45° and an additional 20 cuts were made. The remaining cells were detached by scraping the well with the cell lifter. The cell clumps were broken up by pipetting with a P1000 3-5 times. 500uL of the cell aggregate suspension were transferred to one well of a 6-well plate previously rinsed with Anti- Adherence Rinsing Solution (STEMCELL Technologies, 07010) containing 2.5mL of N2B27 media. Cell aggregates were fed every two days with N2B27 medium until Day 20. Aggregates were fed by transferring all the contents of a well into a 15mL conical tube.

Aggregates were allowed to sink to the bottom of the tube (approximately 3-5 minutes) and two-thirds of the medium were aspirated. To prevent aggregates from clumping, they were gently pipetted 3 times with a P1000. Aggregates were added back to the well with fresh N2B27 medium. On Day 20, aggregates were fed with PDGF medium. PDGF medium was composed of basal media with 1X N2 supplement, 1X B27 minus vitamin A supplement, 10 ng/ml PDGF-AA Protein (R&D Systems, 221-AA), 10 ng/ml IGF-I Protein (R&D Systems, 291-G1-200), 5 ng/ml HGF Protein (PeproTech, 315-23), 10 ng/ml NT-3 Protein (PeproTech, 450-03), 60 ng/ml 3,3′,5-Triiodo-L-thyronine (Sigma-Aldrich, T28), 100 ng/ml Biotin (Sigma-Aldrich, B4639), 1 µM cAMP (Sigma-Aldrich, D0627), and 25 µg/mL insulin. On Day 30, round aggregates with a dark center were selected and seeded on a plate coated with poly-l-ornithine and laminin. Cells were fed every two days by gently removing the two-thirds of the medium with a P1000 and replenishing with fresh medium using a P1000. Cells were fed until Day 65 with PGDF medium.

### MACS to isolate oligodendrocyte progenitor cells

On Day 65 of oligodendrocytes differentiation, MACS was used to isolate OPCs. Cells and aggregates were dissociated by incubating with 2mL Accutase at 37 °C for 30 min. After the incubation, a p1000 was used to mechanically dissociate the cells by gently pipetting 7-10 times. Following mechanical dissociation, accutase was diluted 1:1 with DMEM/F12 supplemented with 10 µM Y27632. Cell suspension was centrifuged at 200× g for 4 min at RT. Media was aspirated and cells were resuspended in DMEM/F12 supplemented with 100ug/mL DNase (Worthington Biochemical, LS002006). After incubating in DNase, cell suspension was further dissociated by gently pipetting 7-10 times. Cells were passed through a 40 µm strainer. Flow-through was collected and centrifuged at 200× g for 4 min at RT. Media was aspirated and cells were resuspended in 120 µL of cold 0.5% BSA made in 1X PBS. Cell suspension was incubated for 10 minutes in the refrigerator.

40µL of the Anti-A2B5 MicroBeads (Miltenyi Biotec, 130-093-392) were added to the cell suspension and and incubated for 15 minutes in the refrigerator. Cells were washed by adding 1mL of 0.5% BSA and spun down at 200g for 4 mins at RT. Supernatant was aspirated completely and resuspend in 500 µL of 0.5% BSA. Cell suspension was applied into MS Columns (Miltenyi Biotec, 130-042-201) attached to MiniMACS™ (Miltenyi Biotec, 130-042-102). Columns were washed with 1 mL of 0.5% BSA three times. Column was removed from the separator and placed it on a suitable collection tube. To wash out cells from the column, 1 mL of 0.5% BSA was added and immediately flushed out the magnetically labeled cells by firmly and slowly pushing the plunger into the column to collect sorted OPCs. Sorted OPCs were seeded on a plate coated with poly-l-ornithine and laminin.

### S63845 treatment of oligodendrocyte progenitor cells

Following sort, OPCs were treated with 2 µM S63845 (MedKoo, 406849) diluted in PDGF media for 48 hours. DMSO (Sigma-Aldrich, D2650) was used as vehicle control. Media with S63845 was replenished at 24 hours.

### Motor neuron differentiation

Motor neuron differentiation protocol was adapted from Ziller and Ortega, 2018^69^. WTC11 human induced pluripotent stem cells (hiPSCs) were dissociated using Accutase and incubated at 37 °C for 5 min. Accutase was diluted with DMEM/F12 medium. A cell lifter was used to remove cells from the well and fully dissociate by pipetting mixture 2–5 times with a p1000 pipette. Cells were centrifuged at 200× g for 4 min at RT. Cells were plated at 8 × 10^4^ cells per well on a Matrigel-coated 6-well plate in E8 supplemented 10 µM Y27632.The following day, E8 was replenished to remove Y27632. Medium changes were performed daily until 1 day following replating when cells reached 80-90% confluency. At this point, differentiation was started (Day 0) by feeding cells with motor neuron induction medium (MNIM) composed of 50% DMEM/F12, 50% Neurobasal media (Thermo Fisher Scientific, 21103049), 1X Penicillin-Streptomycin, 1X MEM Non-Essential Amino Acid, 1X GlutaMAX, 1X N2 supplement, and 1X B27 minus vitamin A supplement. MNIM was supplemented with 10 µM SB431542, 100 nM LDN193189, 1 µM SAG, and 1 µM Retinoic acid. Media was changed daily and SB431542, LDN193189, SAG, and Retinoic acid were added fresh each day. On Day 6, cells were transitioned to MNIM supplemented with 4 µM SU5402 (DNSK International, 215542-92-3), 5 µM DAPT (DNSK International, 208255-80-5), 1 µM SAG, and 1 µM Retinoic acid. Media was changed daily until day 13 and SU5402, DAPT, SAG, and Retinoic acid were added fresh each day. On day 14, neurons were dissociated using Accutase and incubated at 37 °C for 10 min. Accutase was diluted with DMEM/F12 medium. A cell lifter was used to remove cells from the well and fully dissociate by pipetting mixture 2–5 times with a p1000 pipette. Cells were centrifuged at 200× g for 4 min at RT. Cells were plated at 1 × 10^6^ cells per well on a poly-l-ornithine/laminin-coated 6-well plate in motor neuron maturation media (MNMM) supplemented 10 µM Y27632. MNMM was composed of Neurobasal media, 1X Penicillin-Streptomycin, 1X MEM Non-Essential Amino Acid, 1X GlutaMAX, 1X N2 supplement, 1X B27 minus vitamin A supplement, 0.2 ug/mL Ascorbic Acid (Sigma-Aldrich, A4403), 10 ng/mL CNTF (R&D Systems, 257-NT-010), 10 ng/mL BDNF (Peprotech, 11166-BD-010), and 10 ng/mL GDNF (R&D Systems, 11166-BD-010). The following day, MNMM was replenished to remove Y27632. Cells were fed every two days until day 22. To prepare neurons for co-culture, cells replated at 1 × 10^5^ per mL on poly-l-ornithine/laminin-coated plates in MNMM supplemented 10 µM Y27632 on day 23. The following day, MNMM was replenished to remove Y27632. On day 26, motor neurons were transitioned to 50% MNMM and 50% co-culture media. Co-culture media was composed of Neurobasal media, 1X Penicillin-Streptomycin, 1X MEM Non-Essential Amino Acid, 1X GlutaMAX, 1X N2 supplement, 1X B27 minus vitamin A supplement, 20 ug/mL Ascorbic Acid, 10 ng/mL CNTF, 10 ng/mL BDNF, 10 ng/mL GDNF, 60 ng/ml T3, 100 ng/ml biotin, 1 µM cAMP, and 25 µg/ml Insulin.

### Co-culture maintenance

Following S63845 treatment, OPCs were replated at 1 × 10^5^ per mL onto motor neurons. Co-culture was maintained in co-culture media by feeding cells every other day until day 20.

### Western blot

Cells were harvested in lysis buffer composed of PhosSTOP (Roche, 04906837001), PIC (Roche, 04693116001), and PMSF (RPI, P20270-10.0) diluted in 1% Triton (Sigma-Aldrich, T9284). Cells were sonicated to ensure complete cell lysis. 15ug of protein were loaded and ran in 4–20% Mini-Protean TGX precast protein gels (Bio-Rad, 4561094) in Tris-Gly-SDS buffer. Gels were transferred onto polyvinylidene difluoride membranes (Bio-Rad, 1620177) at 4°C overnight. Membrane was blocked in 5% milk (RPI, M17200500) diluted in TBST for one hour at room temperature on a benchtop rocker. Primary antibodies were incubated at 4°C overnight on benchtop rocker. HRP-conjugated secondary antibodies against mouse or rabbit IgG were incubated for one hour at room temperature on benchtop rocker. Blots were developed with ECL Plus Reagent (Thermo Fisher Scientific, 32106) and image on Chemiluminescent Imager. Bands were quantified with Image Studio Lite.

### RNA extraction and cDNA synthesis

Cells were harvested in TRIzol Reagent (Invitrogen, 15596018). RNA isolation was performed following the Invitrogen TRIzol Reagent Protocol. RNA was incubated with DNase I (New England Biolabs, B0303S) at 37 °C for 10 minutes. Reaction was inactivated by adding 0.1 M EDTA (EMD Millipore, 324506) and incubated for 10 min at 75 °C. 10 μL of this volume was used with the High-Capacity cDNA Reverse Transcription Kit (Thermo-Fisher Scientific, 4368814) to obtain cDNA.

### Quantitative RT PCR

1 μg of cDNA sample was used to run RT-qPCR on QuantStudio 3 Real-Time PCR machine with SYBR Green master mix (Thermo Fisher Scientific, 4309155). Manufacturer’s instructions were used to set up the assay.

### Immunofluorescence

Cells were fixed with 4% paraformaldehyde (Electron Microscopy Sciences, 15710-S) in 1X PBS for 30 minutes at room temperature. Cells were incubated in blocking and permeabilization solution composed of 0.3% Triton (Sigma- Aldrich, T9284) and 5% normal donkey serum (Jackson ImmunoResearch, 017-000-121) in 1X TBS for one hour at room temperature. Primary antibodies were diluted in blocking and permeabilization solution and incubated at 4°C overnight. Cells were washed three times with 1X TBS followed by incubation in secondary antibody diluted in blocking and permeabilization solution for one hour at room temperature. Cells were washed three times with 1X TBS. Cells were incubated with Hoechst (1 μg/mL) (Thermo Fisher Scientific, H3570) for 10 minutes. Cells were washed once with 1X TBS and sealed with Fluoromount-G slide mounting medium (Thermo Fisher Scientific, 00-4958-02). Cells were imaged on the Nikon W1 Spinning Disk Confocal 100X Plan Apo 1.49 NA Oil objective and a Prime 95B CMOS camera. Quantification of microscopy images was performed in NIS-Elements (Nikon) or Imaris (Oxford Instruments). Surface Surface Contact Area and Filament Sholl Analysis Imaris XTensions were used for images analyses.

### Statistical analysis

All experiments were performed with a minimum of three biological replicates. Statistical significance was determined by Student’s t-test or one-way ANOVA as appropriate for each experiment. Significance was assessed using Fisher’s protected least significance difference test. GraphPad Prism v10 was used for all statistical analysis (SAS Institute: Cary, NC, USA) and data visualization. Error bars in all graphs represent standard error of the mean unless otherwise noted in the figure legend. No outliers were removed from the analyses. For all statistical analyses, a significant difference was accepted when P < 0.05.

### Antibodies

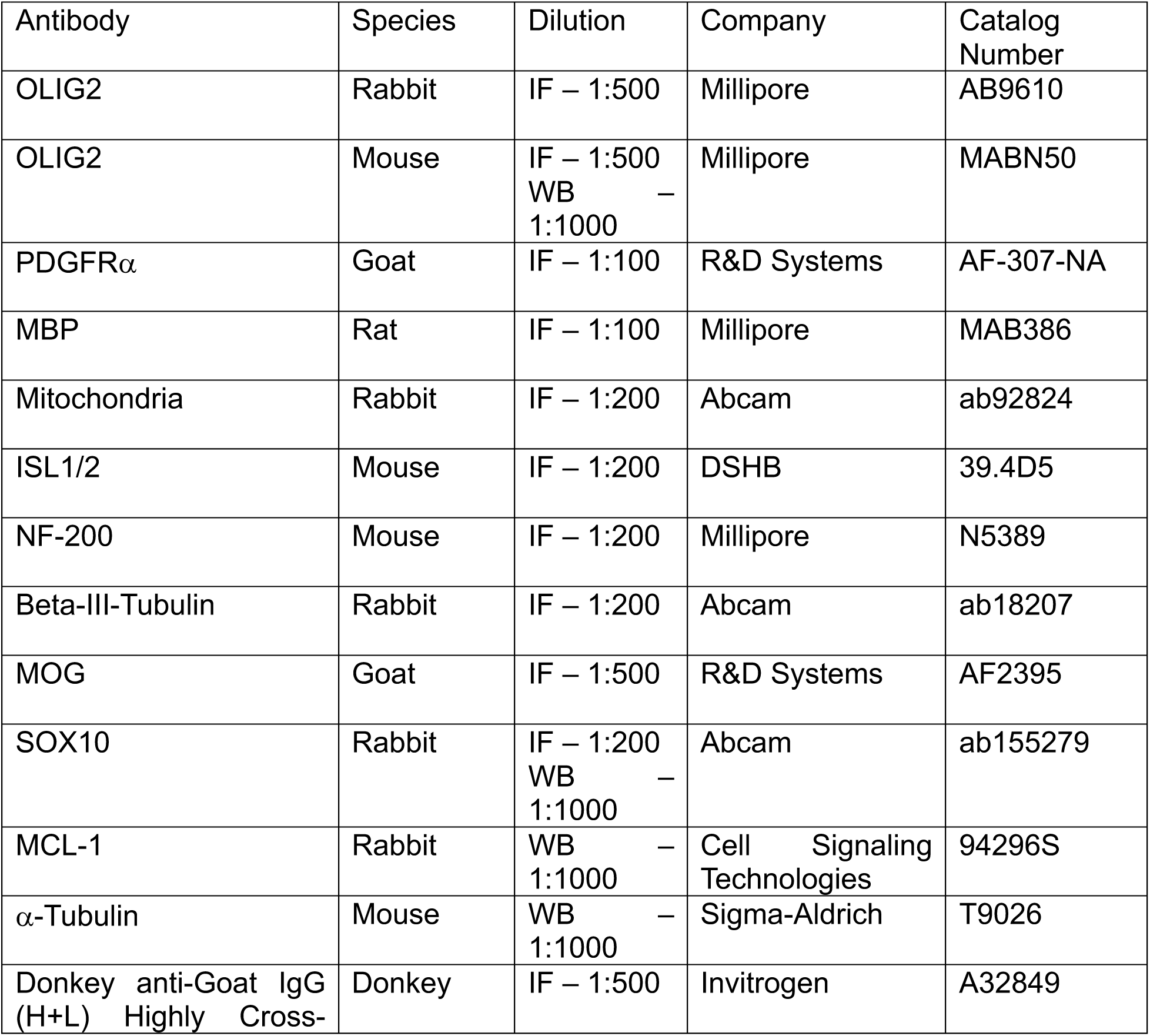
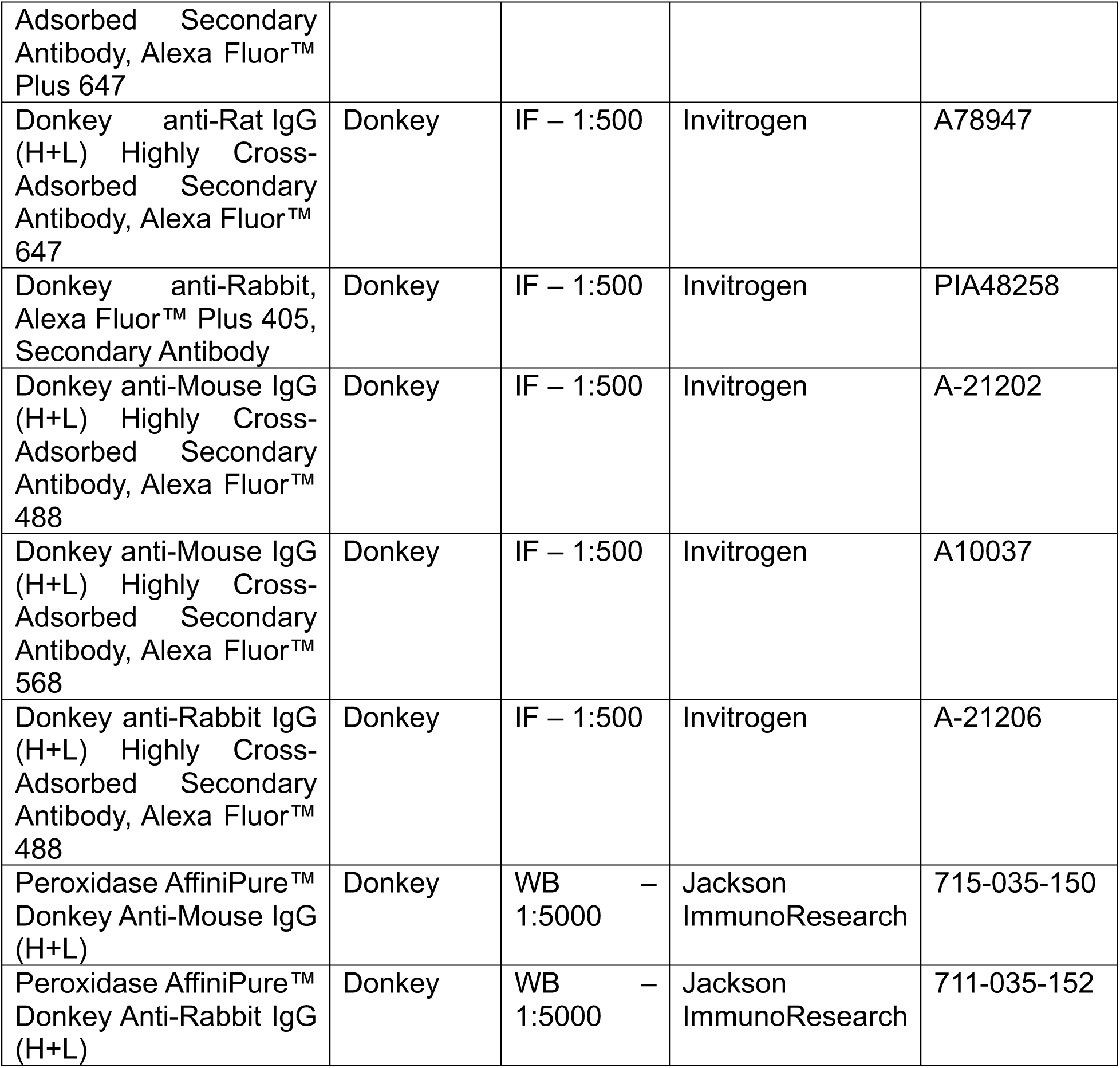

### qPCR primers

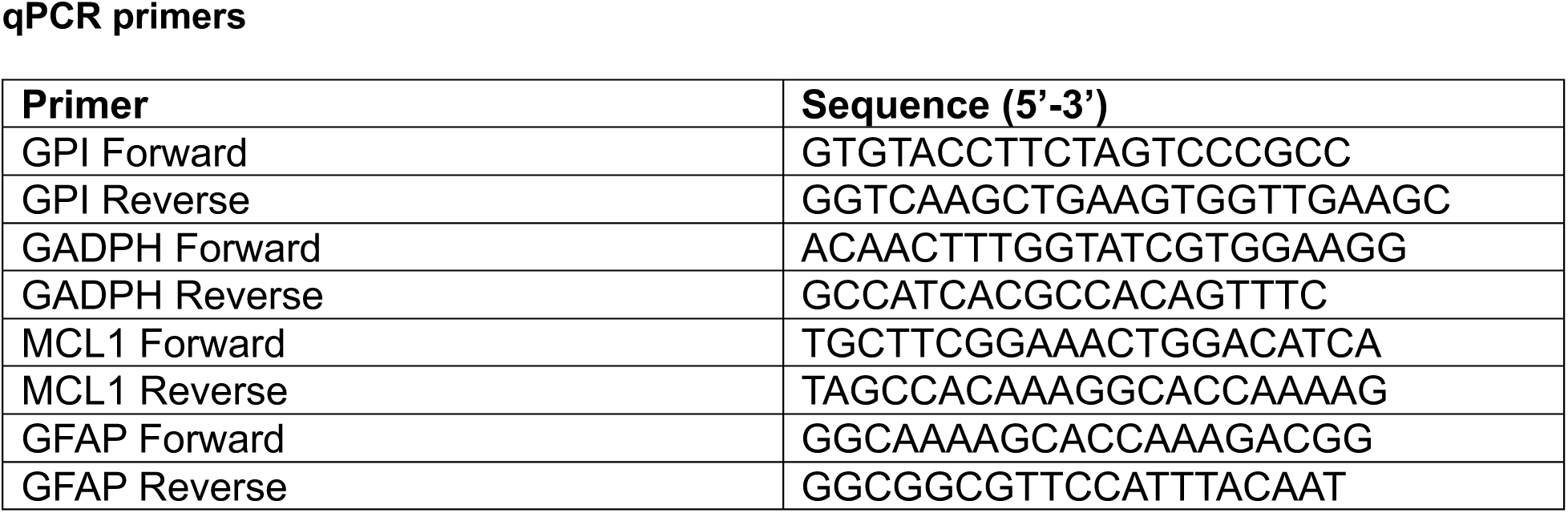
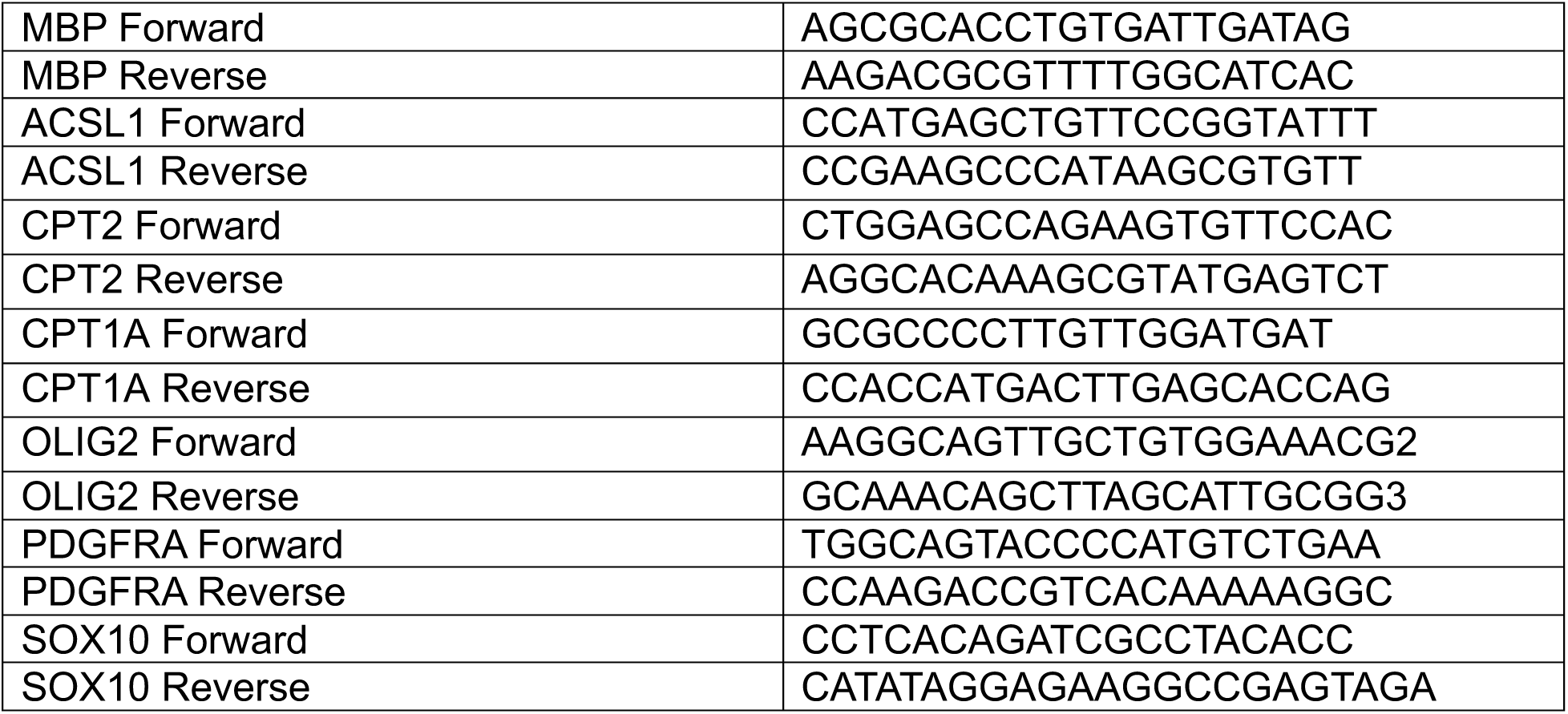

## References

1. Fünfschilling, U. et al. Glycolytic oligodendrocytes maintain myelin and long-term axonal integrity. Nature 485, 517–521 (2012).

2. Ziabreva, I. et al. Injury and differentiation following inhibition of mitochondrial respiratory chain complex IV in rat oligodendrocytes. Glia 58, 1827–1837 (2010).

3. Bame, X. & Hill, R. A. Mitochondrial network reorganization and transient expansion during oligodendrocyte generation. Nat. Commun. 15, 6979 (2024).

4. Knobloch, M. et al. A Fatty Acid Oxidation-Dependent Metabolic Shift Regulates Adult Neural Stem Cell Activity. Cell Rep. 20, 2144 (2017).

5. Asadollahi, E. et al. Oligodendroglial fatty acid metabolism as a central nervous system energy reserve. Nat. Neurosci. 1–11 (2024) doi:10.1038/s41593-024-01749-6.

6. Brinkmann, K. et al. Relative importance of MCL-1’s Anti-Apoptotic versus Non-Apoptotic Functions in vivo. 2023.08.14.553217 Preprint at 10.1101/2023.08.14.553217 (2023).

7. Rasmussen, M. L. et al. A Non-apoptotic Function of MCL-1 in Promoting Pluripotency and Modulating Mitochondrial Dynamics in Stem Cells. Stem Cell Rep. 10, 684–692 (2018).

8. Rasmussen, M. L. et al. MCL-1 InhibitionbySelectiveBH3MimeticsDisrupts Mitochondrial Dynamics Causing Loss of Viability and Functionality of Human Cardiomyocytes. iScience 23, 101015 (2020).

9. Wright, T. et al. Anti-apoptotic MCL-1 promotes long-chain fatty acid oxidation through interaction with ACSL1. Mol. Cell 84, 1338–1353.e8 (2024).

10. Kotschy, A. et al. The MCL1 inhibitor S63845 is tolerable and effective in diverse cancer models. Nature 538, 477–482 (2016).

11. Cleveland, A. H. et al. Oligodendrocytes depend on MCL-1 to prevent spontaneous apoptosis and white matter degeneration. Cell Death Dis. 12, 1133 (2021).

12. Gil, M. & Gama, V. Generation of Myelinating Oligodendrocytes from Pluripotent Stem Cells. (2024).

13. Douvaras, P. & Fossati, V. Generation and isolation of oligodendrocyte progenitor cells from human pluripotent stem cells. Nat. Protoc. 10, 1143–1154 (2015).

14. Shamas-Din, A., Kale, J., Leber, B. & Andrews, D. W. Mechanisms of action of Bcl-2 family proteins. Cold Spring Harb. Perspect. Biol. 5, 1–21 (2013).

15. Sancho, M., Leiva, D., Lucendo, E. & Orzáez, M. Understanding MCL1: from cellular function and regulation to pharmacological inhibition. Febs J. 289, 6209–6234 (2022).

16. Fogarty, L. C. et al. Mcl-1 and Bcl-xL are essential for survival of the developing nervous system. Cell Death Differ. 26, 1501–1515 (2019).

17. Arbour, N. et al. Mcl-1 is a key regulator of apoptosis during CNS development and after DNA damage. J. Neurosci. 28, 6068–6078 (2008).

18. Detmer, S. A. & Chan, D. C. Functions and dysfunctions of mitochondrial dynamics. Nat. Rev. Mol. Cell Biol. 8, 870–879 (2007).

19. Liesa, M., Palacín, M. & Zorzano, A. Mitochondrial dynamics in mammalian health and disease. Physiol. Rev. 89, 799–845 (2009).

20. Robertson, G. L. et al. DRP1 mutations associated with EMPF1 encephalopathy alter mitochondrial membrane potential and metabolic programs. J. Cell Sci. 136, jcs260370 (2023).

21. Baum, T. B., Bodnya, C., Costanzo, J. & Gama, V. Patient mutations in DRP1 perturb synaptic maturation of cortical neurons. 2024.08.23.609462 Preprint at 10.1101/2024.08.23.609462 (2024).

22. Salazar, M. P. R. et al. Mitochondrial fission controls astrocyte morphogenesis and organization in the cortex. 2024.10.22.619706 Preprint at 10.1101/2024.10.22.619706 (2024).

23. Benard, G. et al. Mitochondrial bioenergetics and structural network organization. J. Cell Sci. 120, 838–848 (2007).

24. Picard, M., Shirihai, O. S., Gentil, B. J. & Burelle, Y. Mitochondrial morphology transitions and functions: implications for retrograde signaling? Am. J. Physiol. - Regul. Integr. Comp. Physiol. 304, R393–R406 (2013).

25. Soares, R. et al. Lineage-specific changes in mitochondrial properties during neural stem cell differentiation. Life Sci. Alliance 7, (2024).

26. Hall, A., Giese, N. A. & Richardson, W. D. Spinal cord oligodendrocytes develop from ventrally derived progenitor cells that express PDGF alpha-receptors. Development 122, 4085–4094 (1996).

27. Lu, Q. R. et al. Sonic Hedgehog–Regulated Oligodendrocyte Lineage Genes Encoding bHLH Proteins in the Mammalian Central Nervous System. Neuron 25, 317–329 (2000).

28. Fang, L.-P. & Bai, X. Oligodendrocyte precursor cells: the multitaskers in the brain. Pflugers Arch. 475, 1035–1044 (2023).

29. Barres, B. A. & Raff, M. C. Proliferation of oligodendrocyte precursor cells depends on electrical activity in axons. Nature 361, 258–260 (1993).

30. Wang, J., Xiang, H., Lu, Y., Wu, T. & Ji, G. The role and therapeutic implication of CPTs in fatty acid oxidation and cancers progression. Am. J. Cancer Res. 11, 2477–2494 (2021).

31. Houten, S. M. & Wanders, R. J. A. A general introduction to the biochemistry of mitochondrial fatty acid β-oxidation. J. Inherit. Metab. Dis. 33, 469–477 (2010).

32. Talley, J. T. & Mohiuddin, S. S. Biochemistry, Fatty Acid Oxidation. in StatPearls (StatPearls Publishing, Treasure Island (FL), 2024).

33. Koltai, T., Reshkin, S. J., Baltazar, F. & Fliegel, L. Chapter 4 - Lipid metabolism part I: an overview. in Prostate Cancer Metabolism (eds. Koltai, T., Reshkin, S. J., Baltazar, F. & Fliegel, L.) 71–135 (Academic Press, 2021). doi:10.1016/B978-0-323-90528-2.00013-8.

34. Prew, M. S. et al. MCL-1 is a master regulator of cancer dependency on fatty acid oxidation. Cell Rep. 41, 111445 (2022).

35. Rowitch, D. H. Glial specification in the vertebrate neural tube. Nat. Rev. Neurosci. 5, 409– 419 (2004).

36. Takebayashi, H. et al. The Basic Helix-Loop-Helix Factor Olig2 Is Essential for the Development of Motoneuron and Oligodendrocyte Lineages. Curr. Biol. 12, 1157–1163 (2002).

37. Takada, N., Kucenas, S. & Appel, B. Sox10 Is Necessary for Oligodendrocyte Survival Following Axon Wrapping. Glia 58, 996–1006 (2010).

38. Sock, E. & Wegner, M. Using the lineage determinants Olig2 and Sox10 to explore transcriptional regulation of oligodendrocyte development. Dev. Neurobiol. 81, 892–901 (2021).

39. Ravanelli, A. M. & Appel, B. Motor neurons and oligodendrocytes arise from distinct cell lineages by progenitor recruitment. Genes Dev. 29, 2504–2515 (2015).

40. Liu, Z. et al. Induction of oligodendrocyte differentiation by Olig2 and Sox10: Evidence for reciprocal interactions and dosage-dependent mechanisms. Dev. Biol. 302, 683–693 (2007).

41. Küspert, M., Hammer, A., Bösl, M. R. & Wegner, M. Olig2 regulates Sox10 expression in oligodendrocyte precursors through an evolutionary conserved distal enhancer. Nucleic Acids Res. 39, 1280–1293 (2011).

42. Stolt, C. C. et al. Terminal differentiation of myelin-forming oligodendrocytes depends on the transcription factor Sox10. Genes Dev. 16, 165–170 (2002).

43. Han, H., Myllykoski, M., Ruskamo, S., Wang, C. & Kursula, P. Myelin-specific proteins: A structurally diverse group of membrane-interacting molecules. BioFactors 39, 233–241 (2013).

44. Boggs, J. M. Myelin basic protein: a multifunctional protein. Cell. Mol. Life Sci. CMLS 63, 1945–1961 (2006).

45. Raasakka, A. et al. Membrane Association Landscape of Myelin Basic Protein Portrays Formation of the Myelin Major Dense Line. Sci. Rep. 7, 4974 (2017).

46. Weil, M.-T. et al. Loss of Myelin Basic Protein Function Triggers Myelin Breakdown in Models of Demyelinating Diseases. Cell Rep. 16, 314–322 (2016).

47. Absence of the major dense line in myelin of the mutant mouse ‘shiverer’. Neurosci. Lett. 12, 107–112 (1979).

48. Jean Harry, G. & Toews, A. D. Chapter 4 - Myelination, Dysmyelination, and Demyelination. in Handbook of Developmental Neurotoxicology (eds. Slikker, W. & Chang, L. W.) 87–115 (Academic Press, San Diego, 1998). doi:10.1016/B978-012648860-9.50007-8.

49. Yue, T. et al. A Critical Role for Dorsal Progenitors in Cortical Myelination. J. Neurosci. 26, 1275–1280 (2006).

50. Mei, F. et al. Stage-Specific Deletion of Olig2 Conveys Opposing Functions on Differentiation and Maturation of Oligodendrocytes. J. Neurosci. 33, 8454–8462 (2013).

51. Scolding, N. J. et al. Myelin-oligodendrocyte glycoprotein (MOG) is a surface marker of oligodendrocyte maturation. J. Neuroimmunol. 22, 169–176 (1989).

52. Steiner, P. Brain Fuel Utilization in the Developing Brain. Ann. Nutr. Metab. 75, 8–18 (2020).

53. Mergenthaler, P., Lindauer, U., Dienel, G. A. & Meisel, A. Sugar for the brain: the role of glucose in physiological and pathological brain function. Trends Neurosci. 36, 587–597 (2013).

54. Ebert, D., Haller, R. G. & Walton, M. E. Energy Contribution of Octanoate to Intact Rat Brain Metabolism Measured by 13C Nuclear Magnetic Resonance Spectroscopy. J. Neurosci. 23, 5928–5935 (2003).

55. Panov, A., Orynbayeva, Z., Vavilin, V. & Lyakhovich, V. Fatty Acids in Energy Metabolism of the Central Nervous System. BioMed Res. Int. 2014, 472459 (2014).

56. Yin, F. Lipid metabolism and Alzheimer’s disease: clinical evidence, mechanistic link and therapeutic promise. FEBS J. 290, 1420–1453 (2023).

57. Bullón-Vela, M. V., Abete, I., Alfredo Martínez, J. & Angeles Zulet, M. Chapter 6 - Obesity and Nonalcoholic Fatty Liver Disease: Role of Oxidative Stress. in *Obesity* (eds. del Moral, A. M. & Aguilera García, C. M.) 111–133 (Academic Press, 2018). doi:10.1016/B978-0-12-812504-5.00006-4.

58. Houten, S. M. & Wanders, R. J. A. A general introduction to the biochemistry of mitochondrial fatty acid β-oxidation. J. Inherit. Metab. Dis. 33, 469–477 (2010).

59. Adlkofer, K. et al. Hypermyelination and demyelinating peripheral neuropathy in Pmp22- deficient mice. Nat. Genet. 11, 274–280 (1995).

60. Wolf, N. I., ffrench-Constant, C. & van der Knaap, M. S. Hypomyelinating leukodystrophies — unravelling myelin biology. Nat. Rev. Neurol. 17, 88–103 (2021).

61. Mishra, P. & Chan, D. C. Metabolic regulation of mitochondrial dynamics. J. Cell Biol. 212, 379–387 (2016).

62. Glancy, B., Kim, Y., Katti, P. & Willingham, T. B. The Functional Impact of Mitochondrial Structure Across Subcellular Scales. Front. Physiol. 11, 541040 (2020).

63. Kirischuk, S., Neuhaus, J., Verkhratsky, A. & Kettenmann, H. Preferential localization of active mitochondria in process tips of immature retinal oligodendrocytes. Neuroreport 6, 737–741 (1995).

64. Jiang, Y. et al. Electron tomographic analysis reveals ultrastructural features of mitochondrial cristae architecture which reflect energetic state and aging. Sci. Rep. 7, 45474 (2017).

65. McDonnell, E. et al. Lipids Reprogram Metabolism to Become a Major Carbon Source for Histone Acetylation. Cell Rep. 17, 1463–1472 (2016).

66. Martínez-Reyes, I. & Chandel, N. S. Mitochondrial TCA cycle metabolites control physiology and disease. Nat. Commun. 11, 102 (2020).

67. Wang, L., Owusu-Hammond, C., Sievert, D. & Gleeson, J. G. Stem Cell Based Organoid Models of Neurodevelopmental Disorders. Biol. Psychiatry 93, 622–631 (2023).

68. Jhanji, M., York, E. M. & Lizarraga, S. B. The power of human stem cell-based systems in the study of neurodevelopmental disorders. Curr. Opin. Neurobiol. 89, 102916 (2024).

69. Ziller, M. J. et al. Dissecting the Functional Consequences of *De Novo* DNA Methylation Dynamics in Human Motor Neuron Differentiation and Physiology. Cell Stem Cell 22, 559–574.e9 (2018).

