## Supplementary figures and images for "MCL-1 regulates cellular transitions during oligodendrocyte development"

### Supp Figure 1

#### Supplementary Figure 1

A

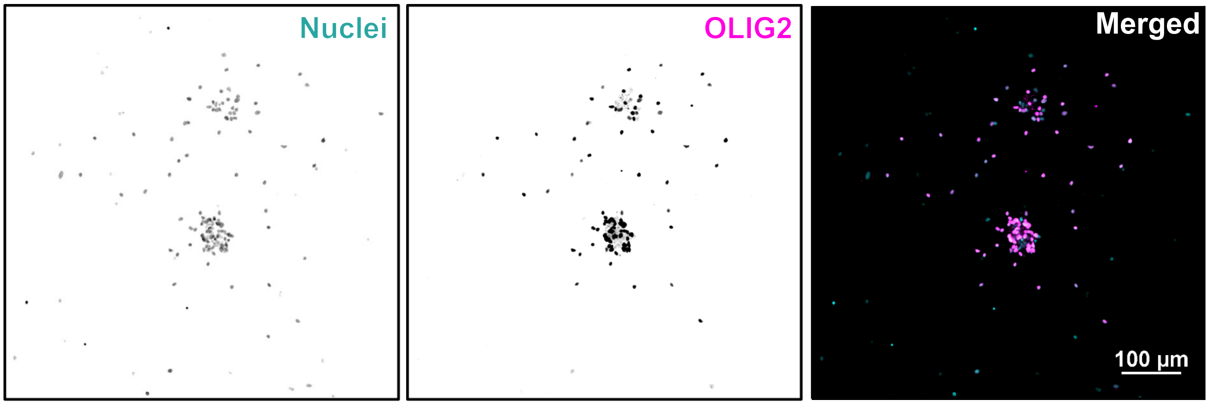


B

| Biological Replicate # | % of OLIG2/Nuclei |
|---|---|
| 1 | 71% |
| 2 | 57% |
| 3 | 80% |
| 4 | 80% |
| Average | 72% |

### Supp Figure 2

#### Supplementary Figure 2

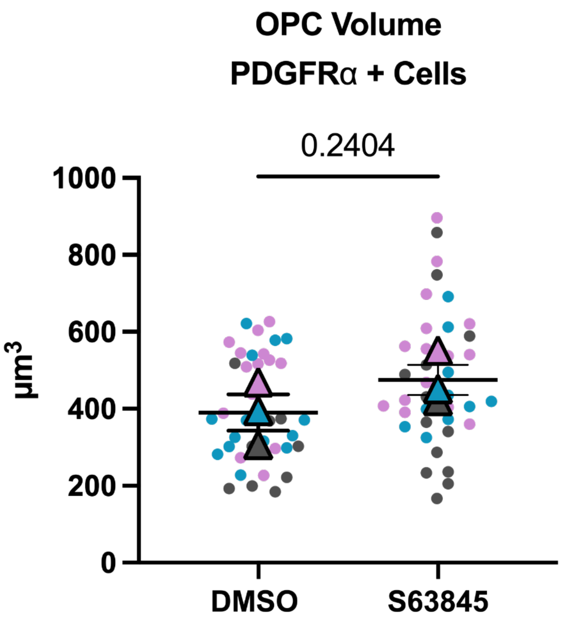

### Supp Figure 3

#### Supplementary Figure 3

A

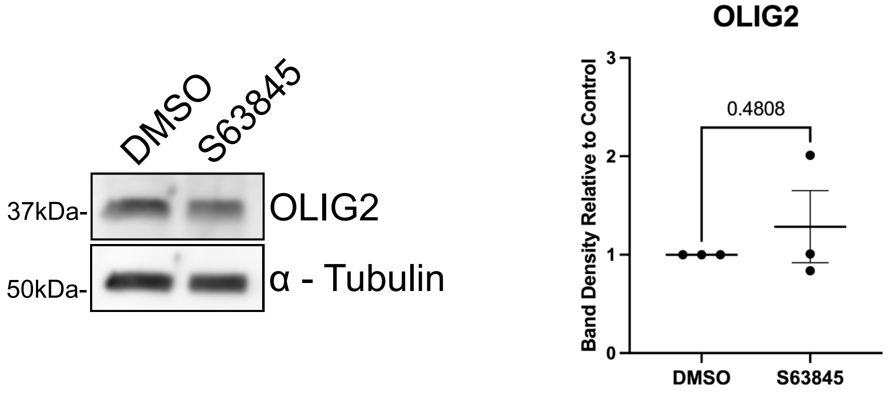


B

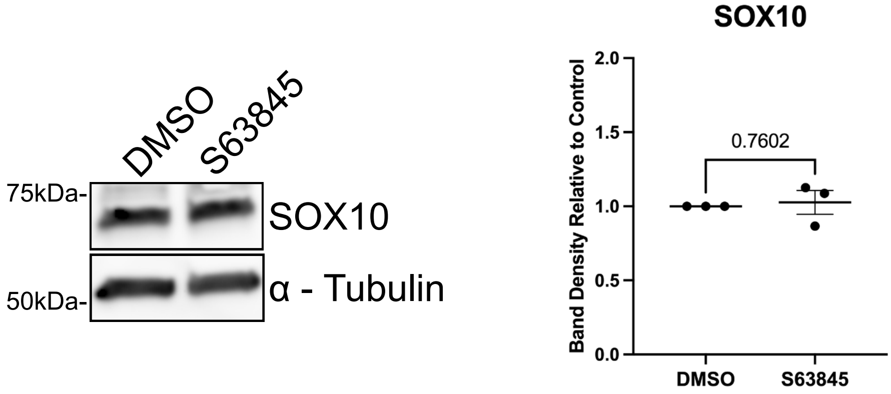

### Supp Figure 4

#### Supplementary Figure 4

A

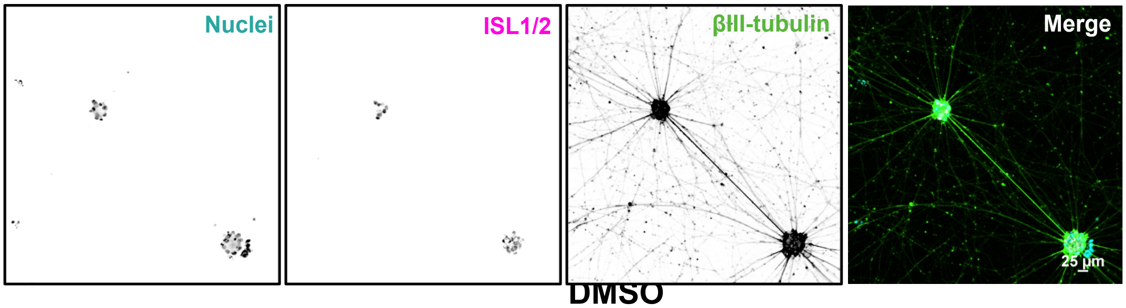


B

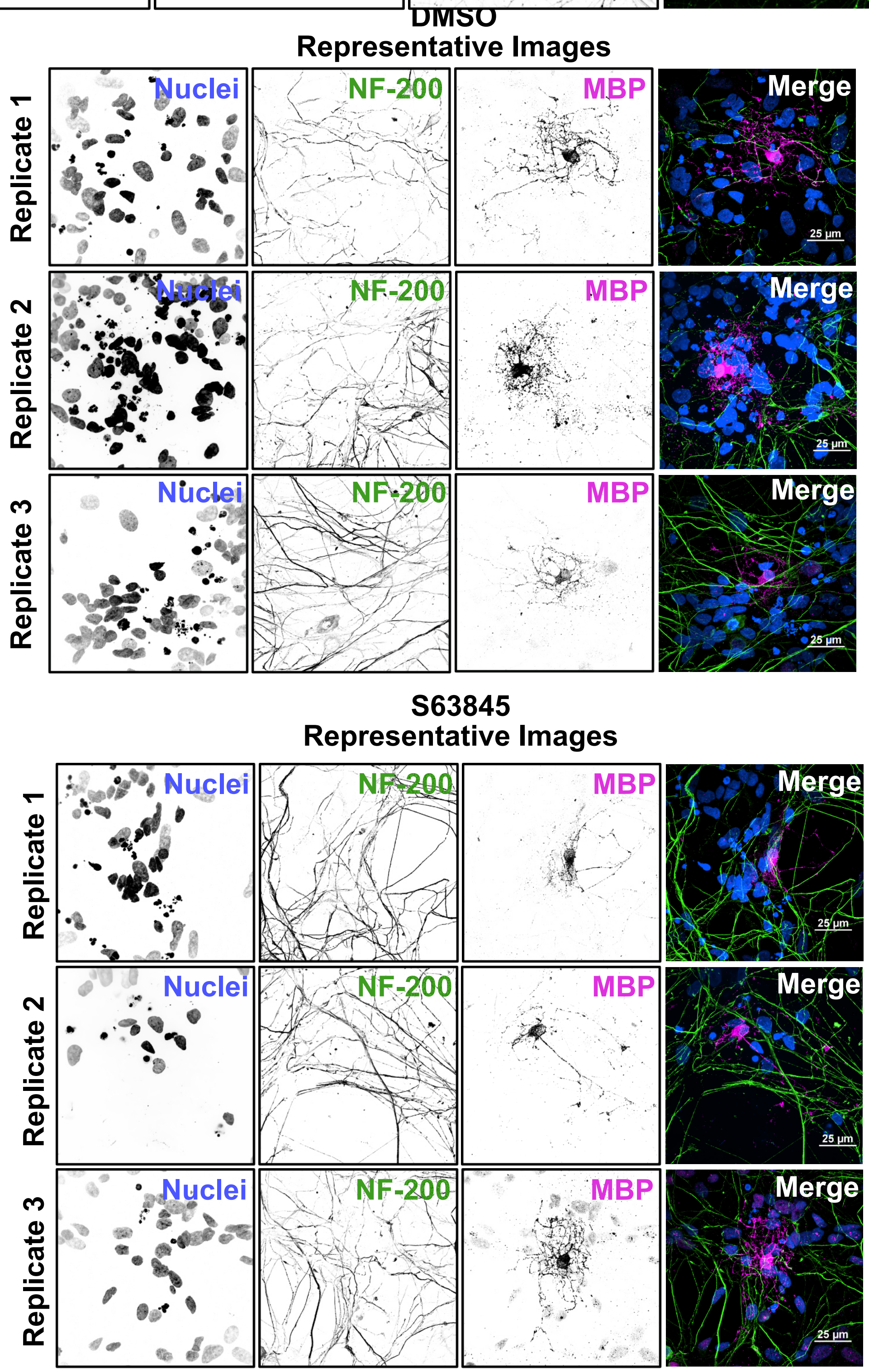

### Supp Figure 5

**Supplementary Figure 5**

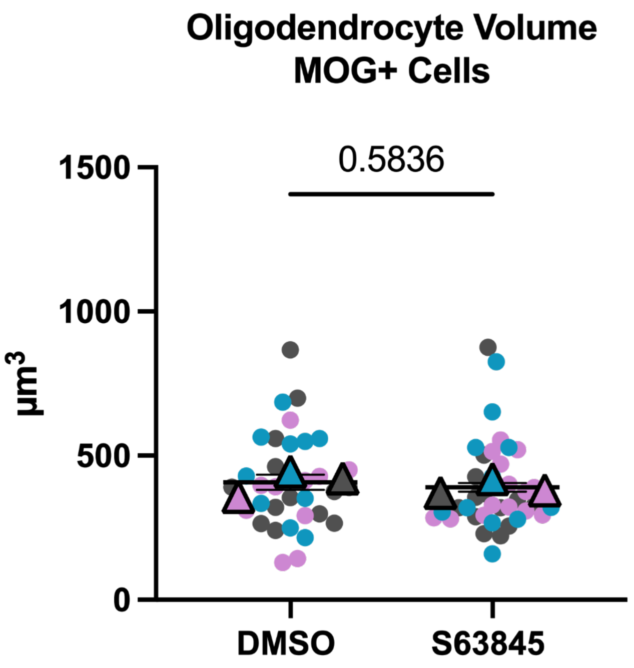
