## Supplementary material for "MCL-1 regulates cellular transitions during oligodendrocyte development": Supp Figure Legends

**Supplementary Figure 1. Validation of oligodendrocyte lineage cell population after cell sort**

(**A**) Spinning disk confocal maximum intensity projections of immunofluorescent staining for nuclei (cyan) and OLIG2 (magenta) in cells after sorting (scale bar = 100μm). (**B**) Chart outlining the percent of OLIG2 positive cells over nuclei in each biological replicate following cell sort.

**Supplementary Figure 2. Volume of PDGFRα positive cells**

(**A**) Volume of platelet-derived growth factor receptor alpha (PDGFRα) positive cells treated with DMSO (vehicle) and S63845. Each color represents a biological replicate (n=3), each dot represents a cell (10-15 per n), each triangle represents mean of biological replicate, analyzed by Student's t-test, error bars represent mean ± SEM. Conditions were blinded to experimenter for 3D reconstructions.

**Supplementary Figure 3. Protein level of OLIG2 and SOX10 in OPCs are not altered following inhibition of MCL-1**

(**A**) Western blot of OLIG2 levels in OPCs following DMSO (vehicle) and S63845 treatment (top panel) and band density quantification (bottom panel). Each dot on graphs represents a biological replicate (n=3), analyzed by students t-test, error bars represent mean ± SEM. (**B**) Western blot of SOX10 levels in OPCs following DMSO (vehicle) and S63845 treatment (top panel) and band density quantification (bottom panel). Each dot on graphs represents a biological replicate (n=3), analyzed by Student's t-test, error bars represent mean ± SEM.

**Supplementary Figure 4. Validation of motor neuron differentiation and representative images of co-culture**

(**A**) Representative spinning disk confocal maximum intensity projections of immunofluorescent staining for nuclei (cyan), ISL1/2 (magenta), and βIII-tubulin (green) in motor neurons derived from human induced pluripotent stem cells (scale bar = 25μm).

(**B**) Representative spinning disk confocal maximum intensity projections of immunofluorescent staining for nuclei (cyan), Neurofilament-200 (NF-200) (green), and myelin basic protein (MBP) (magenta) in day 90 co-cultures with cells that were previously treated as OPCs with DMSO (vehicle) and S63845 (scale bar = 25μm).

**Supplementary Figure 5. Volume of MOG-positive cells**

(**A**) Volume of myelin oligodendrocyte glycoprotein (MOG) positive cells in day 90 co-cultures with cells that were previously treated as OPCs with DMSO (vehicle) and S63845. Each color represents a biological replicate (n=3), each dot represents a cell (10-15 per n), each triangle represents mean of biological replicate, analyzed by Student's t-test, error bars represent mean ± SEM. Conditions were blinded to experimenter for 3D reconstructions.
